# Ranking microbial metabolomic and genomic links in the NPLinker framework using complementary scoring functions

**DOI:** 10.1101/2020.06.12.148205

**Authors:** Grímur Hjörleifsson Eldjárn, Andrew Ramsay, Justin J. J. van der Hooft, Katherine R. Duncan, Sylvia Soldatou, Juho Rousu, Rónán Daly, Joe Wandy, Simon Rogers

## Abstract

Specialised metabolites from microbial sources are well-known for their wide range of biomedical applications, particularly as antibiotics. When mining paired genomic and metabolomic data sets for novel specialised metabolites, establishing links between Biosynthetic Gene Clusters (BGCs) and metabolites represents a promising way of finding such novel chemistry. However, due to the lack of detailed biosynthetic knowledge for the majority of predicted BGCs, and the large number of possible combinations, this is not a simple task. This problem is becoming ever more pressing with the increased availability of paired omics data sets. Current tools are not effective at identifying valid links automatically, and manual verification is a considerable bottleneck in natural product research.

We demonstrate that using multiple link-scoring functions together makes it easier to prioritise true links relative to others. Based on standardising a commonly used score, we introduce a new, more effective score, and introduce a novel score using an Input-Output Kernel Regression approach. Finally, we present NPLinker, a software framework to link genomic and metabolomic data. Results are verified using publicly available data sets that include validated links.

**Author summary:** In this article, we introduce NPLinker, a software framework to link genomic and metabolomic data, to link microbial secondary metabolites to their producing genomic regions.

Two of the major approaches for such linking are analysis of the correlation between sets of strains, and analysis of predicted features of the molecules. While these methods are usually used separately, we demonstrate that they are in fact complementary, and show a way to combine them to improve their performance.

We begin by demonstrating a weakness in the most common method of strain correlation analysis, and suggest an improvement. We then introduce a new feature-based analysis method which, unlike most such methods, does not directly depend on the natural prodcut compound class. Finally, we demonstrate that the two are complementary and proceed to combine them into a single scoring function for genomic and metabolomic links, which shows improved performance over either of the individual approaches.

Verification is done using curated databases of genomic and metabolomic data, as well as public data sets of microbial data including verified links.

## Introduction

Microbial specialised metabolism, i.e. microbial production of metabolites not strictly needed for the survival of the organism, has been a rich source of metabolites for a variety of biomedical applications [1]. Recent advances in genomic analysis [2–4] indicate that microorganisms harbour considerable untapped metabolomic potential [5].

Genes responsible for the production of microbial specialised metabolites are usually grouped into Biosynthetic Gene Clusters (BGCs), contiguous regions of adjacent genes that, taken together, encode enzymes for the production of one or several structurally related metabolites. Several tools exist to predict BGCs from microbial genomes ([4, 6], arguably the most popular of which is antiSMASH [2]. Applying these predictive tools to novel genomes many new BGCs are predicted, the vast majority of which encode unknnown products.

When searching for the products of BGCs, microbial extracts are often profiled using tandem mass spectrometry (MS/MS or MS2). Using this technique, an MS2 spectrum is recorded for a subset of the ionizable metabolites in a culture extract representing the fragmentation of the metabolite, providing useful information as to its structural identity. Data from multiple strains can be combined allowing researchers to see in which strains (or under which growth conditions) particular spectra appear.

Identifying from within these rich metabolomic datasets the particular molecular product (or products) of a BGC is a challenging, but important, problem. Firstly, since rediscovery of known metabolites is a persistent problem in metabolomics [7], linking a spectrum to a BGC with a known metabolite product can help identify the spectrum as belonging to that metabolite, and thus complement database matching as a dereplication strategy. An example of this approach can be seen in the characterisation of a polyketide antibiotic with an elemental formula of C_35_H_56_O_13_ [8]. Secondly, since looking for metabolites similar to a known metabolite is a common starting point in microbial metabolomics (e.g. [9]), establishing the spectrum corresponding to the product of a BGC that is similar to a BGC with a known metabolite product, can be considered particularly useful. This strategy, for example, was used to isolate the novel lanthipeptide tikitericin from a thermophilic bacterium [10]. Lastly, knowing which BGC produces a particular metabolite can give valuable insight for synthetic biology applications, e.g. heterologous expression. This approach was used to identify the normally cryptic BGC responsible for the production of scleric acid, a secondary metabolite with moderate antimicrobial and antitumor activity [11].

Linking BGCs to their metabolite products has largely been done manually and on a small scale, working with single (or small numbers of) strains, either based on similarity to known, existing links (e.g. [10]), or predictions of unique identifying features of the spectrum from the BGCs (e.g. [12]). However, to increase the chance of success, this is increasingly being done at a large scale through the simultaneous analysis of a large number of related strains. The increased throughput makes manual methods prohibitively labour intensive, making development of computational methods for this purpose necessary.

To date, relatively few computational tools exist to aid in this BGC-MS2 linking problem. The problem is challenging: given a collection of BGCs in a strain, and a collection of metabolites produced by the same strain, any given metabolite could, a priori, be produced by any of the BGCs, yielding a huge number of potential links. Compuational approaches have therefore concenrated on scoring possible links, and using the resulting ranking to prioritise links for further analysis. This can be both theoretical (by analysing the BGC to predict the properties of the product) and experimental (e.g. heterologous expression) [13–15]. Existing approaches to compute scores can be broadly classified into two categories: *feature-based approaches*, where the set of MS2 spectra is searched for predicted structural features of the putative product of the BGC (e.g. *peptidogenomcs*, [16]), and *correlation-based approaches*, where similarities in sets of strains containing specific BGCs on the one hand, and specific spectra on the other hand, are used to evaluate the links between BGCs and spectra (*metabologenomics*, [15, 17]).

A few tools exist to aid in *feature-based linking*, such as SANDPUMA [18], GNP [19] and RippQUEST [20]. These tools are each designed to be used for only a single product type such as NRPs, PKs or RiPPs, respectively. Further analysis is therefore required to evaluate the links from different tools against each other.

The most popular *correlation-based* approach is described in [17]. Here, BGCs from different strains are clustered into Gene Cluster Families (GCFs) based on similarity-based distances betweeen BGCs [13, 15] with the assumption that BGCs that are close with regards to this distance will produce identical or structurally similar metabolites. The current state-of-the-art tool for microbial BGC clustering is BiG-SCAPE [13]. GCF and spectra pairs are then scored based upon their shared strains. This approach was for instance used by [15] to link tambromycin to its producing BGC. As well as scoring links between GCFs and spectra, this approach can also be used to score links between GCFs and Molecular Families (MFs) produced by spectral clustering pipelines, such as the *molecular networking* pipeline within GNPS [21].

Fig. 1 is a diagram showing the relationship between the various terms discussed above. For detailed reviews of linking microbial specialised metabolites to their producing BGCs, the reader is referred to [22] and [23]. Note that links can be defined between different entities. For example, on the genomic side, one can consider links involving individual BGCs (feature based linking) or GCFs (feature based or correlation based linking). On the metabolomics side, one can use MFs or individual spectra for either feature- or correlation based linking (both of which have been successfully used, e.g. [9] and [15], respectively).

**Fig 1.**
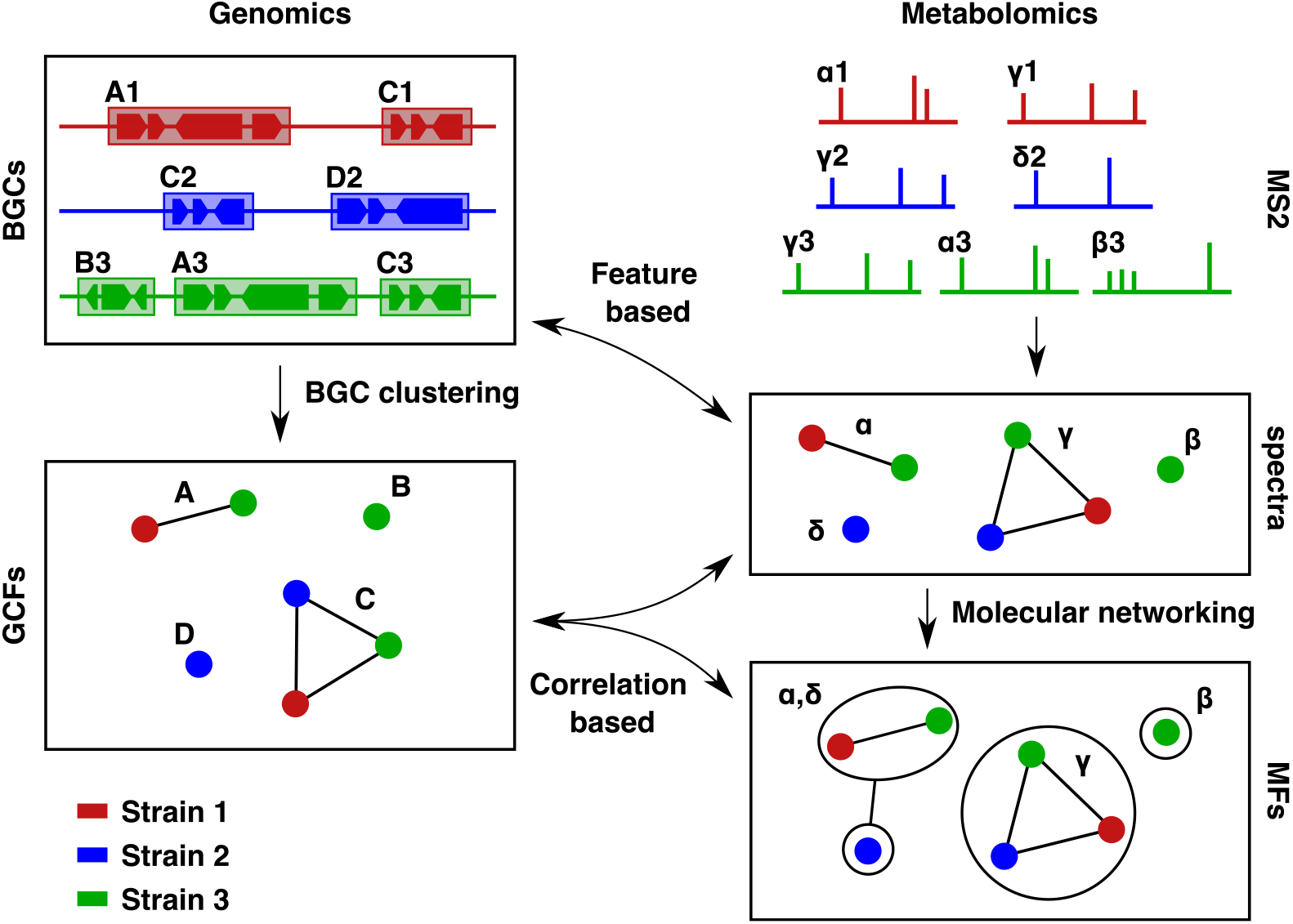
Diagram showing the relationship between the various metabolomic and genomic objects. On the genomics side, BGCs are detected from microbial genomes, colour-coded by strain. These are clustered into GCFs, where each GCF contains BGCs from one or more strains. GCFs can thus also be considered as sets of strains, where each strain contributes at least one BGC to the GCF. On the metabolomics side, MS2 spectra measured in microbial cultures are grouped across strains, so that identical spectra are assigned one or more strains in which they appear. These are further grouped into MFs in a process called Molecular Networking, where each MF consists of one or more related spectra. Both spectra and MFs can likewise be considered as sets of strains where the spectrum, or a spectrum in the MF, is present in the sample for the strain. Feature-based approaches can be used to link BGCs to individual spectra, while correlation-based approaches can be used to link GCFs to either MFs or spectra, based on the pattern of strain contents.

In this work, we address various limitations of the currently available compuational approaches.

- Firstly, as we will show, the most popular strain correlation score [17] has properties that make it impossible to reliably compare score values across links (even links from the same GCF or MF), severly limiting the scores it produces for ranking links. We propose a method for standardising this score such that link scores become comparable across a whole dataset, or for a particular GCF/MF.
- Secondly, although feature-based methods exist for particular specialised metabolite classes (e.g. [18–20]), there is no general feature-based method for ranking links across all classes. Inspired by progress in the area of in silico metabolite identification, in particular [24], we propose the use of on Input-Output Kernel Regression (IOKR) for this task.
- Thirdly, no previous study has attempted to combine feature- and correlation-based scores, despite the fact that they are likely to be complementary. Here we show that strain correlation and IOKR scores are indeed complementary and present a method for combining their scores into a single value.
- Finally, we introduce an open source extendable software framework – NPLinker – into which our developed methods have been implemented that makes it straightforward to import the various data types and will allow accelerated development of new methods for this problem.

## Materials and methods

### Strain correlation scoring

Consider a population of strains, each with a set of predicted BGCs. The union of those sets constitutes a set of BGCs in the population. Assume also that the BGCs have been clustered into GCFs. Similarly, assume that associated with the population is a set of all MS2 spectra for metabolites produced by the population, clustered into MFs. We are interested in scoring these potential GCF-MF links, in order of how likely each GCF is to produce the metabolite giving rise to each MF.

The most common scoring method for scoring a GCF-MF link in metabologenomics is as follows [17]: starting from zero, for each strain in the population,

- add 10 to the score if the strain produces the metabolite and has a BGC in the GCF,
- subtract 10 from the score if the strain produces the metabolite but does not have a BGC in the GCF,
- add 1 to the score if the strain neither produces the metabolite nor has a BGC in the GCF, and
- leave the score unchanged if the strain has a BGC in the GCF but does not produce the metabolite.

Considering the GCF as a set *G* of strains that contribute a BGC to the GCF, and the MF as a set *M* of strains that produce a metabolite in the MF, the score is therefore defined as a function *σ*_corr_(*M, G*) which depends on the size of *M* (#*M*), the size of *G* (#*G*), and the overlap between the two sets, in addition to a set of weights (here 10, −10, 1 and 0, respectively).

While intuitive and easy to interpret, this scoring function, hereafter referred to as *strain correlation score*, has the major drawback of being highly dependent on both total population size and, with potentially greater impact, the size of the GCF and the number of strains that produce the metabolite. A link between a GCF that contains a BGC from every strain in the data set and a moderately-sized MF can easily outscore a link between a smaller GCF and MF that have perfect strain correspondence, even if the evidence for the latter link being valid is stronger, as shown in Fig. 2 (A). This makes comparison between possible links involving GCFs and MFs of different sizes challenging, even within the same data set.

**Fig 2.**
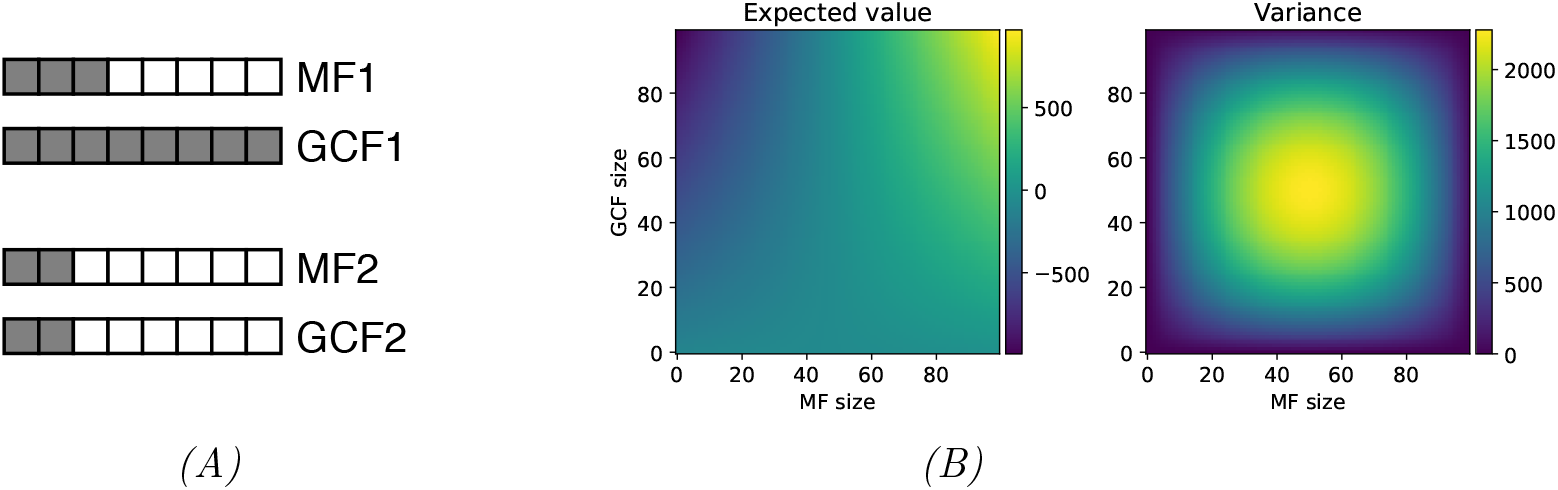
The effect of size on strain correlation scoring. *(A)* Size discrepancy in the strain correlation score for GCFs of varying sizes. Each box represents a strain, with filled boxes denoting that the strain is a member of the GCF or MF, and blank boxes that it is not. The top GCF-MF pair outscores the bottom pair by 30 to 26, despite the bottom pair having arguably stronger correspondence. *(B)* Expected value and variance of the strain correlation score for a population of 100 strains, as a function of GCF and MF sizes. Both the expected value and the variance have a considerable range, rendering comparison between links involving different sizes of GCFs and MFs difficult. For instance, a GCF and MF of size 80 could easily get a score of 500 or higher by chance, while for a GCF and MF of size 20, a score this high would be highly significant.

### Standardisation of the strain correlation score

This effect of GCF and MF sizes on the strain correlation score can be mitigated by standardising the score. For a given GCF and MF, let *G* and *M* be the sets of strains contributing to the GCF and the MF respectively. Assuming the null hypothesis that the strain content of *G* and *M* is not correlated, the probability of a given overlap between the two follows a hypergeometric distribution with the total number of strains as the population size *N*, the size of the molecular family *M* as the number of positives in the population, the size of the GCF *G* as the sample size, and the size of the overlap between the two as the number of positives in the sample.

Letting *m* = #*M*, *g* = #*G*, *o* = #(*G ∩ M*) and *n* = #*N*, we can define the strain correlation 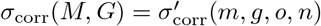 in terms of the sizes of *M*, *G* and the overlap between the two. We can therefore compute the expected value *E*[*σ*_corr_(*M, G*)] as a function of the sizes of *M* and *G* as

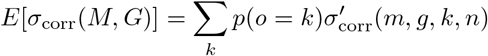

where *k* runs over all possible sizes of #(*M ∩ G*), and *p*(*o* = *k*), the probability of the size of the overlap #(*M ∩ G*) being *k*, follows the hypergeometric distribution as previously stated. The variance can then be computed as

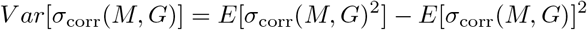

Fig. 2 (B) shows the expected value and variance of the strain correlation score as a function of the sizes of *G* and *M*, for population size #*N* = 100. The expected value varies greatly, especially with the number of strains producing the metabolite (due to the higher magnitude coefficients when the spectrum is present in the strain than when it is absent), and the variation in variance across the domain is also considerable.

We propose to mitigate this by defining a *standardised strain correlation score* for a GCF *G* and MF *M* as

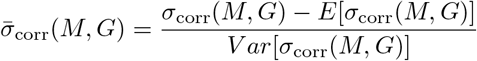

In this version, the score for each prospective link takes into account the sizes of the GCF and the MF and adjusts the score accordingly, making the scores comparable between links involving strain sets of different sizes as is necessary when, for example, comparing scores for different spectra / MF for a particular GCF. In the case of Fig. 2 (A), the standardised scores are 0.0 and 2.65, favoring the bottom pair.

### Input-Output Kernel Regression

If a BGC is known to produces a certain metabolite, the problem of linking a spectrum to that BGC is equivalent to linking a spectrum to the metabolite that the BGC produces. This problem, of matching spectra to molecular structures, is an important problem in metabolomics because it underpins all untargeted metabolomics workflows. This matching is often done in the space of *molecular fingerprints*, which are binary vectors denoting the various properties of the molecule, including the presence or absence of certain substructures. These fingerprints can be derived from candidate structures and effectively predicted from spectra.

Brouard *et al.* propose Input-Output Kernel Regression (IOKR) [24, 25] as a method of ranking a candidate set of chemical structures, given an input spectrum. They compare IOKR with state of the art methods such as CSI:FingerId [26] and demonstrate similar performance with considerably shorter training and classification times, with the training time reduced from 82 hours to under one minute, and classification time reduced by half.

In principle, IOKR works by mapping similarities in the input space (spectra) and the output space (molecular structures) to the molecular fingerprint space and searching a candidate set of metabolites for the closest match in that space. Let 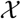 be the space of MS2 spectra and 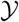 the space of metabolites, with kernel functions 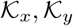, respectively, and 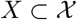 and 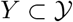 be training sets of paired spectra and metabolites, where each *x_i_* ∈ *X* has a corresponding element *y_i_* ∈ *Y*, Land vice versa. Let 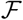 be the space of molecular fingerprints, and 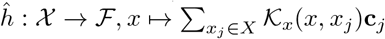, where 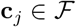 are the fingerprint vectors for the training examples. Finally, let 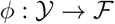 be the mapping expressing the kernel function 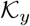 as the inner product in, 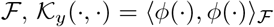. Given a spectrum *x*, IOKR works by searching a set *Y* ^*^ of candidate structures for an element *y* that maximises the expression 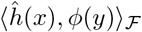. Fig. 3 is an arrow diagram of the IOKR framework.

**Fig 3.**
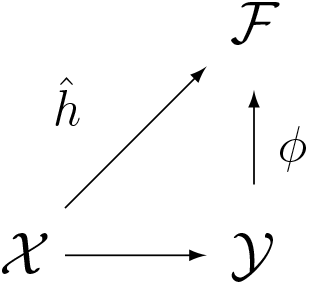
Arrow diagram of the Input-Output Kernel Regression (IOKR) framework. *X* denotes the space of MS2 spectra, 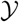 is the space of metabolites, and 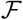 is the shared space of molecular fingerprints. 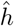 is the (learned) mapping from MS2 to fingerprints, while *ϕ* is the (exact) mapping from metabolites to molecular fingerprints.

For a BGC *g* with an associated molecule 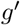 and spectrum *m*, we can then define a link scoring function *σ*_IOKR_ as

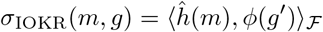

For further details of the underlying mathematics of IOKR, please refer to [25].

### Using IOKR to rank BGC-spectrum links

In the context of linking BGCs and spectra, IOKR can be considered a *feature-based method*. When scoring links between individual BGCs and spectra, however, it does not predict the features of the spectrum directly from the BGC, but uses molecular fingerprints as an intermediate.

The application of IOKR depends heavily on the choice of kernel function on the spectra, and on the choice of molecular fingerprint. For this work, we choose kernel functions that are easy and fast to compute, with the caveat that further optimisation may be possible.

As stated in Section, IOKR works by ranking a set of candidate structures by similarity to a given spectrum. Therefore, this method is not directly applicable to BGCs unless the chemical structure of the metabolite produced by the BGC is known. Generally, this is not the case. However, by considering only the BGCs which have considerable similarity to BGCs in annotated libraries such as MIBiG, structural predictions can be made for a subset of all predicted BGCs. Potential links between spectra and BGCs belonging to this subset can then be ranked using IOKR.

Molecular fingerprints are extracted from SMILES strings using the Chemistry Development Kit [27]. The fingerprint vector is composed of three concatenated sets of fingerprints: *CDK Substructure*, *PubChem Substructure* and *Klekota-Roth* fingerprints.

Taken together, these cover most of the molecular properties described by the fingerprint used by Brourad *et al.* and result in similar performance [24].

As a denoising step, to avoid time-consuming computation of fragmentation trees for the spectra, we filter the input spectra to include only the peaks found in the training data, before using the *Probability Product Kernel* (PPK) [24, 28].

Since the standardised strain correlation score is defined between MFs and GCFs, and the IOKR score between BGCs and spectra, they are not directly comparable. To be able to use them together, we generalise the IOKR score to GCF-MF links by taking the highest BGC-spectrum pair where the BGC is in the GCF and the spectrum in the MF, and assign that score to the GCF-MF pair, i.e. for a GCF *G* and a MF *M*, and a scoring function *σ*_IOKR_ scoring BGC-spectrum links, we define a second function with the sets of GCFs and MFs as a domain, by setting

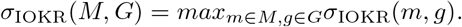

### Combining strain correlation and feature-based scores

So far, computational efforts to find GCF-MF links have mostly been based on either the *feature-based* or the *correlation-based* approach. However, both approaches have limitations. For example, correlation-based methods have trouble prioritising singleton links within the same strain, i.e. when the same strain is associated with multiple singleton GCFs and MFs. Feature-based methods on the other hand rely on being able to predict distinguishing features that can be detected in MS2 spectra from the BGCs, and since the same features are often present in multiple GCFs and MFs, this can yield multiple potential links with the same score.

Because the two approaches are based on very different principles, they are likely to be at least in part complementary, with each one sensitive to things that the other is not. We demonstrate that combining the two approaches by looking at potential links that are highly ranked using both increases the ratio of previously validated links to all links in the joint top percentiles compared to using either one of the approaches.

### NPLinker

To facilitate analysis of paired genomics and metabolomics data sets, we developed NPLinker, a Python module to accelerate and support the process of automatically linking GCFs or their BGCs with observed mass spectra. NPLinker accepts genomic outputs from antiSMASH and BiG-SCAPE (including reference BGCs from the MIBiG database [29]), and metabolomic output from the public, community-driven Global Natural Products Social (GNPS) knowledge base [30]. Additionally, it includes integration with the *Paired omics Data Platform* [31] to retrieve paired public genomics and metabolomics data (https://pairedomicsdata.bioinformatics.nl).

In addition to the framework, NPLinker includes a user interface for visual inspection of the potential links, and can be used as a stand-alone web app. For ease of use, it can be run in a Docker container, working with either local data, links to data sets in the Paired omics Data Platform or a mixture of both.

After loading the metabolomic and genomic data, including automated running of BiG-SCAPE if required, links can be sorted, inspected and filtered by various scoring functions or combinations thereof, and visualised either in form of tables or graphically as networks of connected strains. A screen shot and further documentation for NPLinker are included in Supplementary Information.

NPLinker creates objects for spectra, MFs, BGCs and GCFs in the data set, maintaining the hierarchical relationship between them, and keeps track of strain ID or IDs associated with each object, as well as strain aliases. Various scoring functions can be used to evaluate the links between metabolomic and genomic objects, both scoring functions that are supplied with NPLinker and custom scoring functions. Objects can also be filtered by various criteria such as strain association, inclusion in MIBiG, and annotations. Fig. 4 shows how NPLinker combines the various data sources.

**Fig 4.**
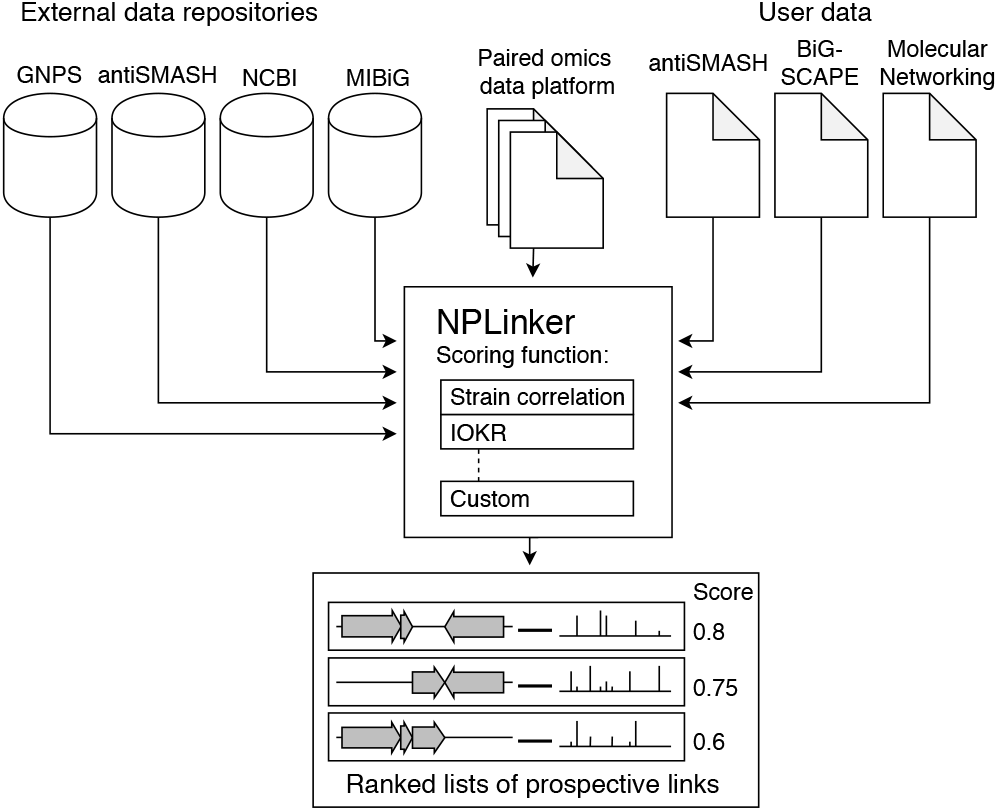
Diagram of the NPLinker module. The NPLinker module helps with automatically linking GCFs and MFs. It integrates metabolomic and genomic data sets, using either external sources, user-provided data, or a mixture of both, and ranks potential links between metabolomic and genomic objects by given scoring functions, either built-in or user-defined.

### IOKR training data

The IOKR model is trained on a set of spectrum-molecular fingerprint pairs. Since molecular fingerprints can be computed from structural annotations, such as SMILES [32] or InChI-strings [33], we use library MS2 spectra from the public, community-driven GNPS knowledge base [30] as a training set for the IOKR model. We use the same training data set as Brouard *et al.* [24], which consists of 4138 spectra from GNPS with structural annotations. The spectra are for metabolites from a variety of sources, including microbial, plant and human metabolites.

## Test data

### Pairing the MIBiG and GNPS databases

In recent years, the MIBiG database [29] has emerged as a central repository of characterised microbial BGCs. The current version, 2.0 [34], contains close to 2000 BGCs, most of which have structural annotations. Many tools, including antiSMASH and BiG-SCAPE, use MIBiG as part of their analysis to quantify similarity of unknown to known BGCs.

In particular, antiSMASH can be configured to compute the similarity of detected BGCs to MIBiG entries and return for each detected BGC a (ranked) list of similar BGCs in MIBiG, using the *known cluster blast* feature [2]. Assuming that similar BGCs give rise to similar compounds, we used this list in turn to assign one or more molecular structures to BGCs, according to how many high-scoring matches are found in MIBiG (or none, if no match is found).

In its current form, MIBiG has no information about the MS2 spectra of the metabolites produced by the BGCs. However, the entries are annotated with structures, so the structural annotations included in GNPS can be used to link GNPS spectra to their corresponding MIBiG BGCs. In this way, we built a set of known BGC-spectrum pairs. To avoid distinguishing between metabolites based on properties absent from an MS2 spectrum, e.g. the chirality of the metabolite, this linking is done using only the first part of the InChI-key of each metabolite. This yields 2966 BGC-spectrum pairs, each with an associated metabolite, which can be used to evaluate the IOKR model proposed in this paper. These pairs include 2069 unique spectra and 242 unique MIBiG BGCs. This list is included in the Supplementary Information.

### Validated links from published data

A major problem in the development of methods to link BGCs to spectra for collections of microbial strains is the lack of ground truth data. Given a data set, consisting of a collection of GCFs for a population of strains, and a collection of MFs for the same strains, the number of potential GCF-MF links is vast. Because microbial secondary metabolism is largely controlled by BGCs, many of these potential links will be true, in that the BGCs in the GCF are responsible for the production of the molecules in the MF. Only a very small number of these true links have been verified, however. Any effective scoring function will therefore have a large number of true but unverified links towards the top end of the distribution.

The performance of a scoring function can be measured by the ratio of true links to all links at the top end of the distribution. The ideal scoring function would assign all true links a higher score than all false links, and as any improvement in ranking should increase the portion of true links and decrease the portion of false links towards the top of the distribution, comparing two scoring functions can be done by considering this ratio for both functions. While the set of verified links for each data set constitutes only a very small subset of the actual links, the same principle can be used to compare scoring functions using only the verified subset of links, i.e. comparing the ratio of *verified* links to all links, instead of the ratio of *true* links to all links, as increasing the ratio of true links to all links would in particular increase the ratio of verified links to all links.

Recognising the difficulty of obtaining ground truth data in this field, Schorn *et al.* recently developed the Paired Omics Data Platform documenting the location of genomic and metabolomic data sets from microbial experiments, with a focus on data sets with BGCs and MS2 spectra [31]. This gives a repository of established links in various data sets, citing the articles in which the links were verified, which can then be used to evaluate scoring methods for prospective links between BGCs and spectra, in terms of relative over-representation of established links towards the upper end of the distribution of scores. From this platform we concentrated on three data sets each with numerous validated links: MSV000078836 [35], MSV000085018 [36] and MSV000085038 [37], hereafter referred to as Cruüsemann, Gross and Leao, respectively.

The Cruüsemann data set consists of 120 microbial strains with 8 established links between a BGC and a MF, the Gross data set consists of 7 strains with 9 established links between a BGC and a MF, and the Leao data set contains 4 strains with 5 established links between a BGC and a specific MS2 spectrum. After downloading the strain assemblies and metabolomics data, the genomes were run through antiSMASH v5.0.0 for BGC detection and BiG-SCAPE v1.0.0 to cluster the BGCs into GCFs. The established links are of various product types, although primarily polyketides and NRPs, with sizes ranging from 7000 to 20000 nucleotides, or between four and 102 genes. A full list of BGCs, their product types and sizes is in supplementary materials.

The Paired Omics Data Platform (PoDP) links are represented by molecular families or MS2 spectra in specific GNPS data sets, and by MIBiG IDs in specific strains. To map the BGC links back to the genomes in question we used antiSMASH to score the correspondence between the MIBiG entries and the detected BGCs, returning for each BGC a (possibly empty) list of MIBiG matches, along with a score for each match. This means that a verified link from the PoDP can link a single spectrum to multiple GCFs, all of which we consider as verified. This can indicate potential splitting of a cluster of similar BGCs, or ambiguity in product type for the BGCs.

## Results

### Standardising the strain correlation score

As the Crüsemann data set contained the largest number of strains of the three data sets being considered, it was selected to evaluate the standardised strain correlation score. To do this, we examine the distribution of scores for validated links in relation to the scores for all hypothetical links. The more effective a scoring method is, the higher it should rank the correct GCF-MF links compared to the rest of the links. Therefore, the mean score for validated links should be higher than the mean score for all potential links, so the null hypothesis for testing the validity of the scoring function is that both distributions of scores (for validated links and all links) have the same mean.

For the raw strain correlation score, the mean score is 83.514 for all links, and 14.667 for validated links, i.e. a lower mean score for the validated links than for all links, which is the opposite of what would be expected. Standardising the score gives a mean score of −0.006 for all links, and 3.672 for validated links (Table 1). The distributions of the scores of the validated links amongst all links are shown in Fig. 5 for both the standardised and raw versions of the score. Similar trends can be observed in the Leao and Gross data sets, see Supplementary Information.

**Table 1.**
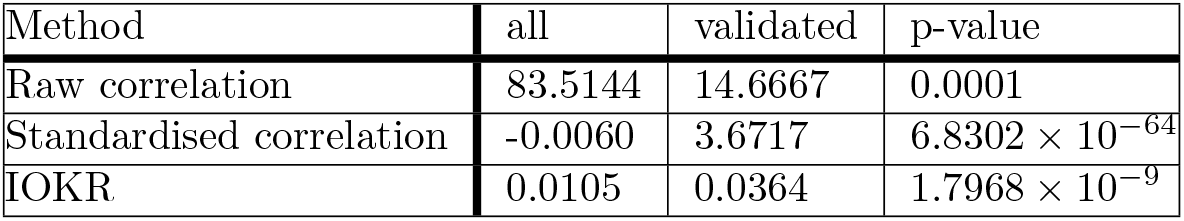
Mean scores for all links and the subset of validated links in the Cruüsemann data set.

**Fig 5.**
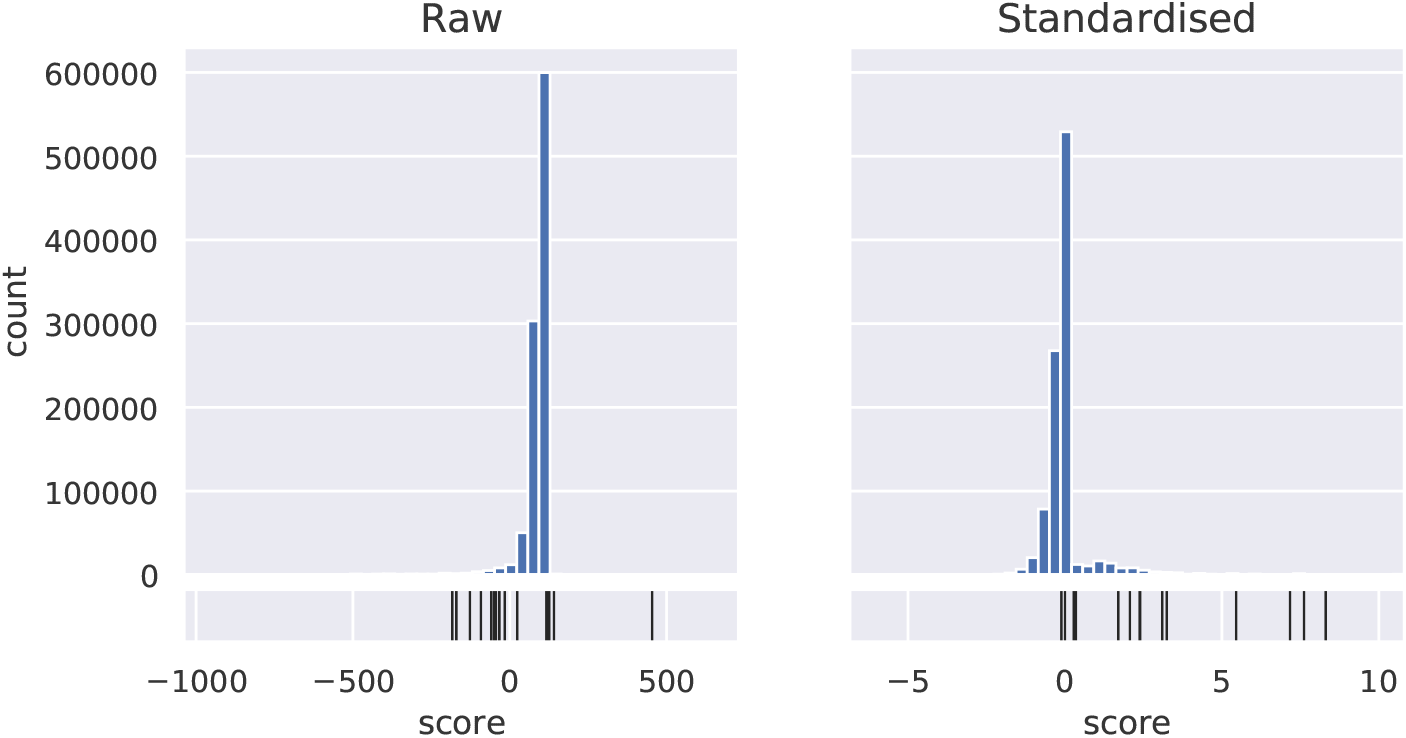
Distribution of validated links among scores. Distribution of the raw and standardised strain correlation scores, as well as the distribution of the scores for validated links (in black) relative to the distribution of scores for all links, in the Cruüsemann data set. The standardised score has a more pronounced tail at the top end, which includes 13 out of 15 validated links, whereas many of the validated links score relatively low on the distribution of the raw scores.

Rows two and three of Table 2 show the proportion of verified links among the top scoring links for the raw and standardised correlation scores. In the Cruüsemann data set, where the strain correlation scoring is most relevant, the proportion is considerably higher for the standardised correlation score relative to the raw correlation score.

**Table 2.** Proportion of verified links among all possible GCF-MF links in the three data sets.

| Data set | Crüsemann | Gross | Leao |
| --- | --- | --- | --- |
| Total | $1.50 \times 10^{-5}$ | $9.96 \times 10^{-6}$ | 0.00086 |
| Top raw corr. | $6.22 \times 10^{-5}$ | $3.48 \times 10^{-5}$ | 0.00266 |
| Top std. corr. | 0.00013 | $7.69 \times 10^{-5}$ | 0.00385 |
| Top IOKR | $6.00 \times 10^{-5}$ | $1.99 \times 10^{-5}$ | 0.00107 |
| Top comb. | **0.00046** | **0.00019** | **0.00680** |

The results demonstrate the importance of our proposed standardisation process.

### Evaluating the IOKR scoring function using MIBiG

As stated earlier, the training set used to build the IOKR model includes metabolites from sources other than microbial. As the performance evaluation of IOKR does not break down performance by metabolite sources, to evaluate the performance of IOKR on microbial specialised metabolites specifically, we tested the method on the paired MIBiG/GNPS data by matching each spectrum to the candidate set consisting of all structures associated with an MIBiG entry. For each spectrum, IOKR returns an ordered list of all metabolites in the candidate set. As multiple BGCs can produce the same metabolite, and as the same BGC can be associated with a number of structurally related metabolites, yielding multiple BGC-MS2 links for the same metabolite.

To translate the ranking of the metabolites into the rank of a BGC among a set of BGCs, we consider the highest-ranking metabolite that the BGC produces. Each metabolite that scores higher than the correct metabolite is associated with one or more BGC. The rank of the correct BGC is the number of distinct BGCs associated with a metabolite that scores higher than the correct metabolite.

For comparison purposes, a baseline score was estimated by randomising the rank of the structures for each spectrum, and the same process was repeated to assign a BGC using the randomised score.

Table 3 shows the top-*n* performance of IOKR, i.e. how often the ‘true’ BGC match for a given spectrum is among the top *n* matches returned by IOKR, for a selection of *n*. IOKR outperforms the baseline by a considerable margin, especially at low values of *n*, with an AUC (i.e. the area under the top-*n* curve for varying values of *n*) of 0.6534 compared to 0.5209 for the null distribution. In this context, the AUC can be considered as a measure of how close we are to the top hit being the correct one in all instances (AUC of 1.0) compared to a randomised baseline.

**Table 3.** Top-***n*** accuracy, and AUC, of IOKR on MIBiG data.

|  | top-1 | top-5 | top-10 | top-20 | top-200 | AUC |
| --- | --- | --- | --- | --- | --- | --- |
| data | 0.1208 | 0.1708 | 0.1870 | 0.2121 | 0.2946 | 0.6534 |
| random | 0.0 | 0.0014 | 0.0044 | 0.0103 | 0.1486 | 0.5209 |

### Evaluating the performance of IOKR

Similarly to the evaluation of the standardised strain correlation score, we can observe the distribution of the scores for the validated links among the scores for all potential links in the Cruüsemann data set. Out of 3316 BGCs in the data set, 2242 could be assigned structure based on similarity to MIBiG entries, and used as candidate set for the 6246 MS2 spectra in the data set. As can be seen from Fig. 6, the upper end of the distribution for the IOKR score contains a relatively high proportion of the validated links, with the mean score of 0.0105 for all links and 0.0364 for validated links (Table 1). Results for other data sets can be found in Supplementary Information.

**Fig 6.**
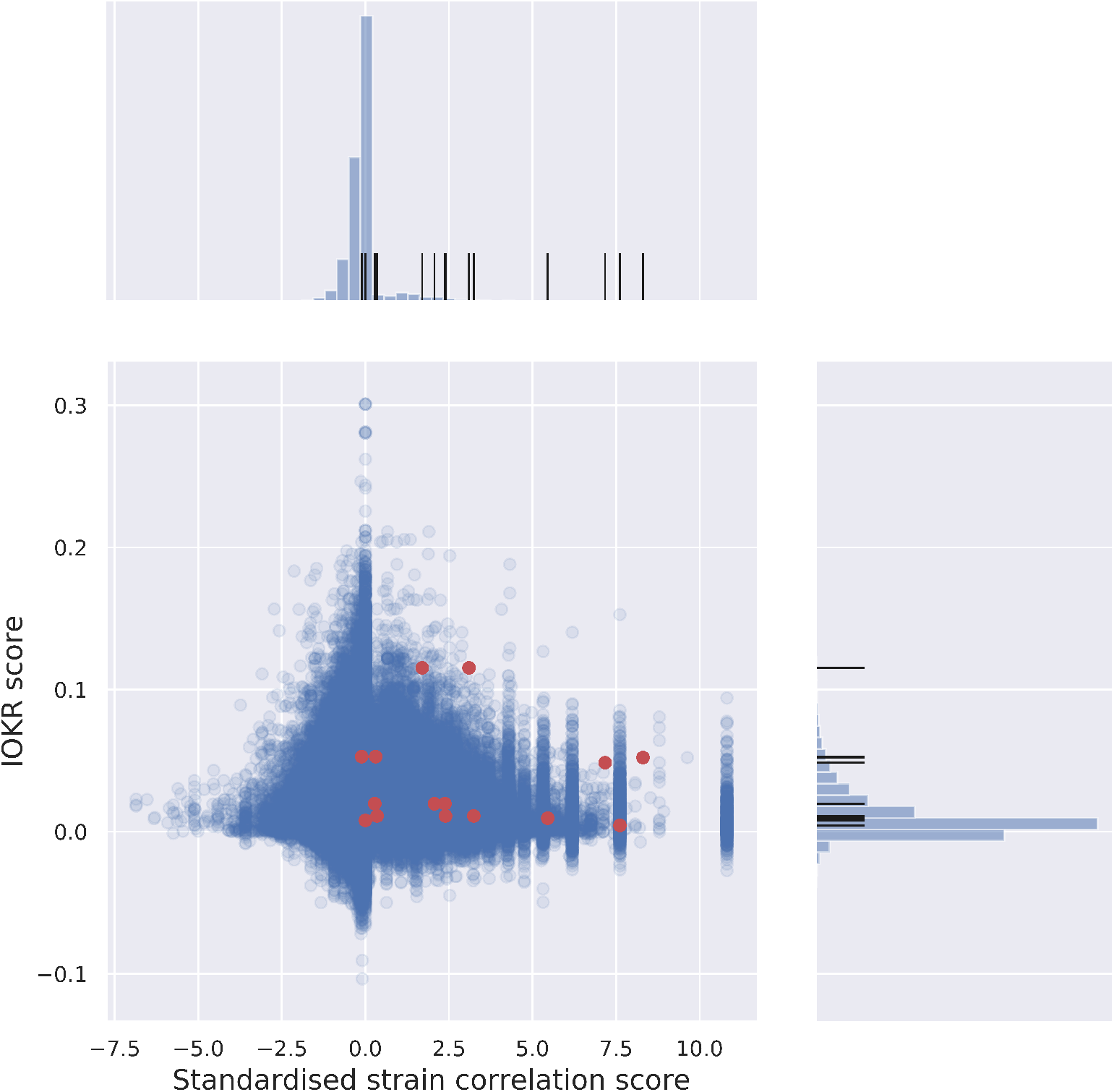
Correlation of IOKR- and strain correlation scores. IOKR- and strain correlation scores for all potential links in the Cruüsemann data set, with histograms of the scores. Verified links are marked on the joint plot, and marked on the distribution histograms. Verified links are concentrated in the upper-right quadrant, i.e. score relatively high on both axes. See Supplementary Information for results for the two further data sets.

### Complementarity of IOKR and strain correlation scoring

If multiple scoring methods are complementary then using them in concert makes sense. One way to demonstrate the efficacy of individual scores is to test whether or not the upper percentiles of their distributions are enriched with verified links, i.e. contain a relatively higher number of verified links compared to the data set as a whole. Table 2 shows the proportion of links that are validated across the whole set of links (*Total*; top row), and for links above the 90th percentile for raw correlation, standardised correlation and IOKR scores (second, third and fourth rows resp.). We can see that for all methods, the 90th percentile contains a higher proportion of validated links than across all links, and the standardised correlation score consistently improves upon the raw correlation score. Note that the rather low value of proportions throughout is largely due to the small number of verified links. The number of true hits in the dataset (and in the 90th percentiles) is likely to be much higher. The final row in Table 2 shows the proportions when looking above the 90th percentile for both the IOKR and standardised scores together. This gives the highest proportions across all three datasets, demonstrating their complementarity, and hence the potential in combining the scores. The distribution of the scores for the Cruüsemann data set, and the relative score of the verified links, can be seen in the histograms of Fig. 6.

Since the number of verified links in each data set is small (ranging from 5 to 15), we can pool the links across the three datasets to get a clearer sense of the statistical significance. Considering the 90th percentile per data set for both scores, and adding up the numbers of links in each category (verified or unverified, and scoring above 90th percentile for either or both scores) for all the data sets, both the IOKR and standardised strain correlation scores are significantly enriched (*p*-value of 0.0139 and 2.483×10^−11^, respectively) for validated links. Furthermore, the set of links scoring above the 90th percentile on both scores is significantly enriched compared to the set that exceed either of the individual scores (*p*-value of 2.633×10^−4^ and 0.0208 starting from IOKR and standardised strain correlation scores, respectively). Relevant tables are provided in supplementary materials for 90th and 95th percentiles.

### Scoring potential links for a particular BGC

A common approach to establishing correspondence between a GCF and a metabolite is to start with a GCF with established homology to a particular MIBiG BGC. The question then becomes one of ranking the potential GCF-MF links *for that particular* GCF to find the most likely product. Starting from a GCF, one can order the list of potential GCF-MF links by standardised strain correlation score, and use the IOKR score to further prioritise the ordered list for verification, or vice versa.

Starting with a GCF in the Cruüsemann data set with homology to a particular MIBiG BGC, and a verified link to a MF, Table 4 shows how many out of the 3094 potential links involving that GCF score as high or higher than the verified link on the IOKR score (col. 2), the standardised strain correlation score (col. 3), or both scores (col.7). For instance, 1151 out of 3093 links including BGC0001228 (retimycin A, validated link excluded) have an equal or higher standardised strain correlation score than the correct link, of which 140 also have an equal or higher IOKR score. Similarly, 15 out of 3093 links containing BGC0000241 (lomaiviticin A, validated link excluded) have an equal or higher standardised strain correlation score, none of which has an equal or higher IOKR score than the correct link. The distributions of scores for a selection of the BGCs belonging to validated links can be seen in Fig. 7

**Table 4.** Scoring function performance.

| MiBiG ID | Rank of verified link | | | | | no. of links w. higher $\sigma_{\text{IOKR}}$ and $\bar{\sigma}_{\text{corr}}$ than the validated link |
| --- | --- | --- | --- | --- | --- | --- |
| | $\sigma_{\text{IOKR}}$ | $\bar{\sigma}_{\text{corr}}$ | $\ell_{\frac{1}{2}}$ | $\ell_1$ | $\ell_2$ | |
| *BGC0001228* | 384 | 1152 | **253** | 271 | 317 | 141 |
| *BGC0000241* | 119 | 16 | **5** | 12 | 14 | 1 |
| *BGC0000333* | 1810 | 25 | 32 | **12** | **12** | 12 |
| *BGC0000827* | 6 | **1** | 2 | **1** | **1** | 1 |
| *BGC0001830* | 673 | **7** | **7** | 9 | 24 | 4 |
| *BGC0000137* | 80 | 16 | **2** | 14 | 49 | 1 |
| *BGC0000137* | 78 | 189 | **12** | 29 | 66 | 3 |
| *BGC0000137* | **798** | 1968 | 1232 | 997 | 920 | 534 |
| *BGC0000137* | 636 | 612 | **409** | 642 | 735 | 120 |
| *BGC0000137* | **6** | 287 | 36 | 214 | 261 | 1 |
| *BGC0000137* | **6** | 126 | 14 | 105 | 122 | 1 |
| *BGC0000137* | 6 | 216 | **4** | 85 | 203 | 1 |
| *BGC0000137* | 184 | 282 | **146** | 271 | 285 | 17 |
| *BGC0000137* | 184 | 101 | **28** | 92 | 99 | 7 |
| *BGC0000137* | 184 | 90 | **19** | 89 | 88 | 7 |
| no. of lowest ranks | 3 | 2 | **10** | 2 | 2 |  |

**Fig 7.**
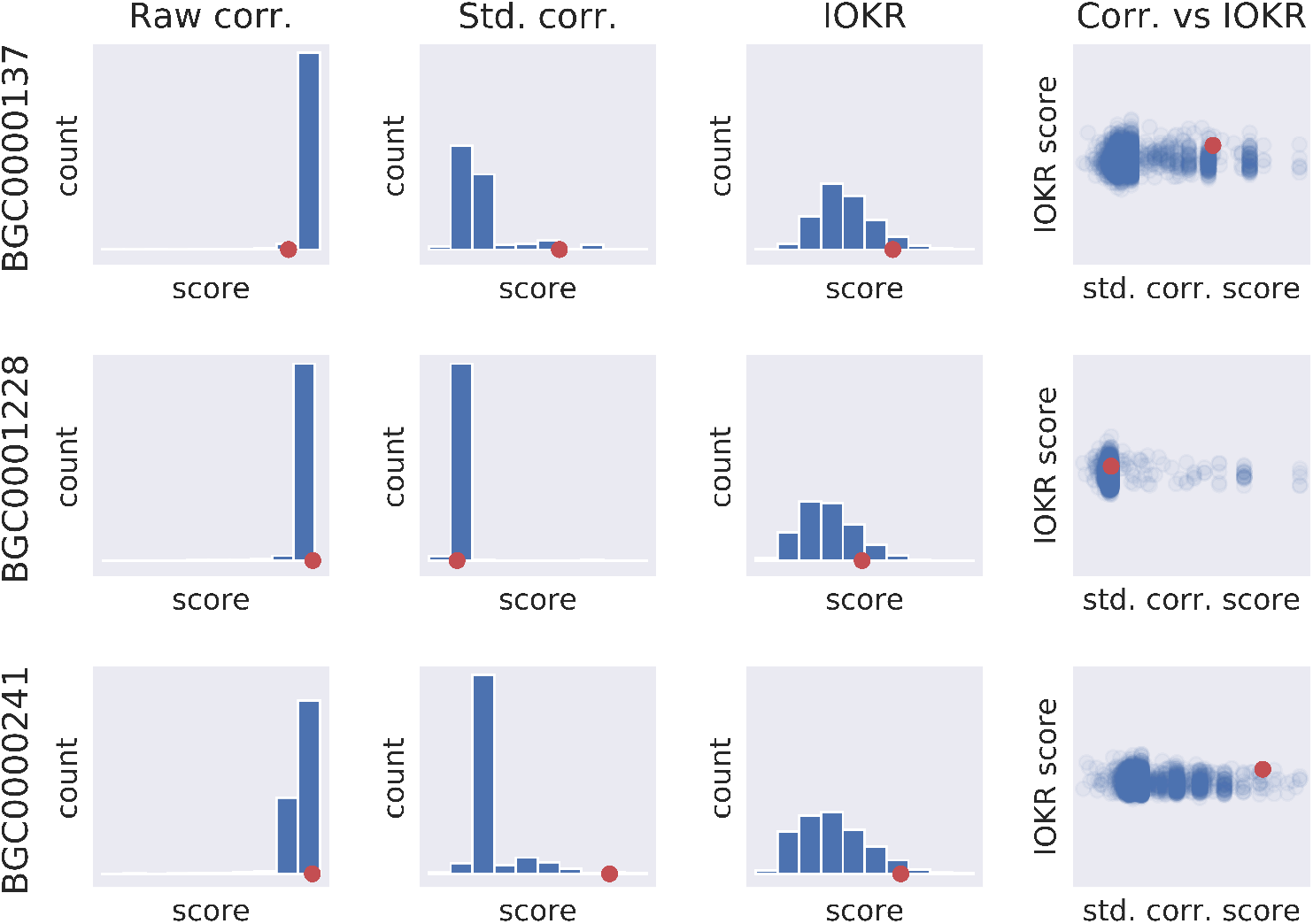
Scores starting from particular GCF. Position of the score for the validated GCF-MF pair within the distribution of the scores of the links between that particular GCF and all MFs, for a selection of established links in the Cruüsemann data set (rows). The first three columns show histograms of the raw and standardised versions of the strain correlation score, as well as the IOKR score, for all links including a given GCF, with the score of the correct link indicated. The last column shows the standardised correlation score (*x*-axis) and IOKR score (*y*-axis) for the same links, again with the correct link indicated. Both IOKR and the standardised correlation scores tend to put established links higher in the distribution of scores for the GCF in consideration, than the raw correlation score. Furthermore, some of the validated links score relatively higher on IOKR than the standardised strain correlation score, and vice versa, suggesting that the two scores complement one another. For full results, as well as for other data sets, please refer to Supplementary Information.

Columns two and three show the rank of the verified link ordered by *σ*_IOKR_ and 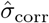. Columns four through six show the rank of the verified link using the *ℓ_p_* scoring functions. The last column shows how many links rank higher than the validated link on both *σ*_IOKR_ and 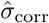. Multiple appearances of the same MIBiG IDs are due to antiSMASH mapping the same ID to multiple GCFs, as discussed earlier.

Whilst this, and the previous section, demonstrates clearly that the scores are complementary, this information in itself does not help us to rank the potential links, since it requires knowledge of the score of the true link, which is not known a priori. One way of using both scores simultaneously to rank this list is to combine them into a single score. Optimising the exact combination of scores is outside the scope of this article, however, we can demonstrate the utility of doing so, with the understanding that further optimisation may be possible.

To make the IOKR score comparable to the standardised correlation score, we can standardise it in a similar manner to the strain correlation score. Letting *σ*_IOKR_ be the IOKR score, we define the standardised IOKR score 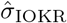 as

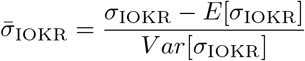

where the variance and expected value are taken over the set of all potential links.

In order to define the function used to combine the two scores, we can consider a two-dimensional space where each axis corresponds to one of the scores, and concentric rings centered at the origin. In the quadrant where both scores are positive, quadrant I, the links further from the origin should be ranked higher. Looking at the set of all circles centered at the origin, we can then order the links by the radius of the circle on which they lie, with a preference for larger radius.

Looking at Fig. 6, however, the scores extend along the axes (close to each score being 0), rather than forming a circle. Therefore, rather than restrict ourselves to computing distance from the origin wih the Euclidean norm, *ℓ*_2_, we can consider the more general *ℓ_p_*-norm where in the 2-dimensional case

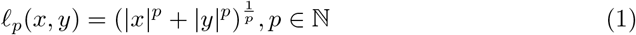

In our application, we can also consider values of *p* such that 0 *< p <* 1. Although in those cases *ℓ_p_* is not a norm, since it does not fulfill the triangle inequality, this is not a problem when using it to rank combined scores. Fig. 8 shows the set of points (*x, y*) such that *ℓ_p_*(*x, y*) = 1 for three values of *p*, demonstrating the parallels of the circle in 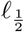 1 to the distribution of scores in Fig. 6.

**Fig 8.**
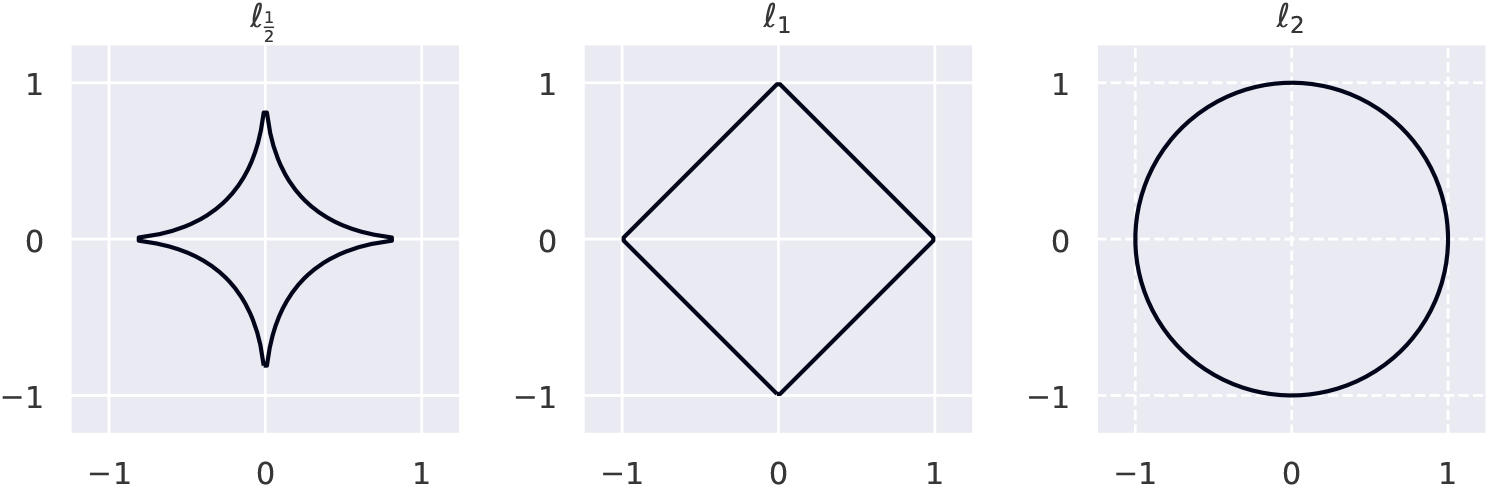
Combining scores. The set of points (*x, y*)such that *ℓ_p_*(*x, y*) = 1, for three different values of *p*.

In the positive quadrant (I) of the coordinate system, we can use Equation 1 directly, with *x* as the standardised correlation score and *y* as the standardised IOKR score. To penalise scores in the other quadrants, particularly quadrant III, we can multiply the *x*- and *y*-terms with the corresponding sign. Using 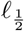, our combined scoring function is therefore

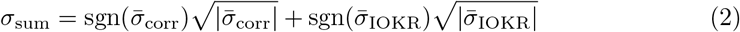

where *σ*_corr_ is the standardised correlation score and *σ*_IOKR_ the standardised IOKR score, and sgn is the sign function mapping positive values to 1 and negative values to −1.

The rank of the verified links according to the various scoring functions can be seen in Table 4. The first two columns show the number of links scoring higher or equal to the verified link ordered by the IOKR and the standardised correlation scores, while the next three columns show the same where links have been ordered by the combined scores for 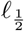, *ℓ*_1_ and *ℓ*_2_. In 10 out of the 15 verified links considered, the 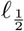 score assigns the highest rank to the verified link, including in three out of the first five cases where the link is unambiguous. For instance, the verified link for BGC0001228 (retimycin A) is ranked at number 253 and for BGC0000241 (lomaiviticin A) at number 5, both of which are considerably higher than for either scoring function on their own, as well as for the other values of *p* tested.

While other values of *p*, or other functions to combine the scores, may improve these results, we believe our analysis demonstrates the utility of using both the standardised strain correlation score and the IOKR score as complementary scoring functions to prioritise the list of potential links.

## Discussion

We have shown that standardising the strain correlation score makes it more effective at prioritising validated links relative to all links, and introduced IOKR as a complementary feature-based scoring function. The major strength of IOKR combined to other feature-based methods is that it is not dependent on product type, but yields scores that are directly comparable between different product types. We have also demonstrated that considering both scoring methods at the same time increases the ratio of validated links among high-scoring links. Furthermore, by pairing the MIBiG and GNPS databases, and using the Paired omics Data Platform, we introduced data sets to test the efficiency of the scoring methods, both separately and combined.

The standardised strain correlation score still suffers from the drawback inherent in correlation-based scoring, of not being able to distinguish between potential links showing the same pattern of strain presence or absence. An obvious example is prioritising multiple singleton GCFs and MFs for the same strain. As a complementary scoring function to the strain correlation score, IOKR does not have this limitation.

While it is not theoretically dependent on product type, due to insufficient test set size, breaking down performance by product type to verify this is currently difficult. A drawback of the current IOKR scoring method is its reliance on MIBiG homology to assign molecular structures to BGCs, which is needed to compute the molecular fingerprint. This restricts its use to those BGCs which show considerable homology with MIBiG entries. While still useful in this form, predicting molecular fingerprints directly from BGCs would broaden the applicability of the scoring function.

As stated earlier, IOKR is also highly dependent on the choice of both kernel function and molecular fingerprints. As the molecular fingerprints denote particular substructures of the molecules, creating additional fingerprints that specifically target molecular substructures seen in secondary metabolites may improve performance. Similarly, the kernels used on the MS2 space can almost surely be further optimised, both through Multiple Kernel Learning [25] and other approaches such as vector embeddings [38].

Leveraging the correspondence between data from multiple microbial strains is an important tool to aid in linking BGCs to spectra. Standardising the strain correlation score by taking into account the number of strains involved in the link makes the score more useful by minimising the effect of strain count on the score, thereby no longer favouring common BGCs and molecules. We also introduced IOKR as a novel feature-based scoring method for potential BGC-spectrum links, and showed how the two methods complement one another by demonstrating the relative enrichment of previously validated BGC-spectrum links amongst the top-scoring links in both scoring functions. By using both scores simultaneously, the prioritisation of hypothetical links can be made more effective. Finally, we introduced the NPLinker framework to aid in prioritising BGC-spectrum links for further research. We believe that our work provides the natural products community with new tools that ease the combined analysis of genome and metabolome mining approaches. This may pave the way toward the discovery of novel chemistry that is much needed to fight off pathogenic bacteria, viruses, and insects.

## Supporting information

**S1 Appendix. Strain correlation score** *p***-value**. Calculating the *p*-value of a given strain correlation score, given the GCF and MF sizes.

**S2 Appendix. The NPLinker framework.** More detailed description of the NPLinker framework.

**S3 Appendix. NPLinker documentation**. Information on documentation of NPLinker structure and functionality.

**S4 Table. Product type of verified links**. A table showing BGC size (in nucleotides and number of genes) and product type for each of the verified links in the data sets.

**S5 Fig. Raw vs. standardised strain correlation scores**. Histograms showing the distribution of raw and standardised strain correlation scores for the data sets, as well as positions of verified links within the distribution.

**S6 Table. Sizes of data sets**. Table showing number of BGCs, number of BGCs with assigned structure and number of spectra per data set.

**S7 Fig. IOKR score of validated links** Histograms of the distribution of IOKR scores for the data sets, as well as positions of verified links within the distribution.

**S8 Table. Number of validated links in higher percentiles**. Number of links (validated vs. total) for the different scoring functions and data sets, in the 90th and 95th percentiles of either or both scores.

**S9 Table. Comparison of score distributions**. Means of scores for validated vs. all links in the data sets.

**S10 Fig. IOKR vs. correlation score**. Figures showing the IOKR- and standardised strain correlation scores for the data sets, with the validated links marked.

**S11 Fig. Score distributions for a particular BGC**. Figures showing the distributions of scores starting from BGCs in validated links.

**S12 File. Linked MIBiG and GNPS databases**. File containing GNPS id, MIBiG id and SMILES string, linking the two databases.

**S13 File. High-scoring links from the Cruüsemann data set**.

**S14 File. High-scoring links from the Leao data set**.

**S15 File. High-scoring links from the Gross data set**.

## Acknowledgments

JJJvdH acknowledges an ASDI grant from the Netherlands eScience Center − NLeSC (grant no. ASDI.2017.030). AR, KRD and SR acknowledge funding from the Biotechnology and Biological Sciences Research Council (BB/R022054/1). KRD, SR and SS are supported by a Carnegie Trust Collaborative Research Grant. JR acknowledges funding from the Academy of Finland (grants 310107 and 334790) and Scottish Informatics and Computing Science Alliance (SICSA) distinguished visiting fellow scheme.

## Notes

### Competing Interest Statement

The authors have declared no competing interest.

### Summary of Updates

Section on combining scores for ranking likns for a particular BGC added. Introduction updated with an explanatory diagram. Current state of the art, and contribution, clarified.

