## Supplementary material for "Ranking microbial metabolomic and genomic links in the NPLinker framework using complementary scoring functions": Suplemental information and data: README_supplementary.txt

Supplementary data with "Ranking microbial genomic and metabolomic links in the NPLinker framework using complementary scoring functions"
by Grimur Hjorleifsson Eldjarn, Andrew Ramsey, Justin J. J. van der Hooft, Katherine R. Duncan, Sylvia Soldatou, Juho Rousu, Rónán Daly, Joe Wandy, and Simon Rogers

gnps_mibig_smiles.csv: The GNPS-MIBiG pairs from matching InChI keys on both data sets.
links_*_99.csv: GCF-MF links scoring above the 99th percentile on both standardised strain correlation and IOKR scores. 
