## Supplementary material for "Ranking microbial metabolomic and genomic links in the NPLinker framework using complementary scoring functions": Suplemental information and data: supplementary_info.pdf

Supplementary information

Grímur Hjörleifsson Eldjárn,
Andrew Ramsay,
Justin J. J. van der Hooft,
Katherine R. Duncan,
Sylvia Soldatou,
Juho Rousu,
Rónán Daly,
Joe Wandy,
and Simon Rogers

September 24, 2020

#### 1 Strain correlation score $p$-value

Consider the population of strains as a set $N$, with cardinality $\#N$, and consider the GCF $G$ and molecular family $M$ as subsets of $N$. A strain correlation scoring function $\sigma$ is a function taking as arguments two subsets $G, M \subseteq N$, representing the GCF and the MF respectively, and assigning to the pair a real-valued score $s$.

The scoring function we use is the scoring function defined in [1], which is defined as

$$\begin{aligned}\sigma(M,G) =& \alpha\#(M\cap G) + \beta\#(N\setminus(M\cup G)) + \gamma\#(G\setminus(M\cap G)) + \delta\#(M\setminus(M\cap G)) & (1)\\ =& \alpha\#(M\cap G) + \beta(\#N - \#(M\cup G)) + \gamma(\#G - \#(M\cap G)) + \delta(\#M - \#(M\cap G)) & (2)\end{aligned}$$

where $\alpha = 10$, $\beta = -10$, $\gamma = 1$ and $\delta = 0$. The coefficients $\alpha$, $\beta$, $\gamma$ and $\delta$ correspond to the strain being in both the GCF and MF sets ($\alpha$), neither of the sets ($\beta$), the GCF set but not the MF set ($\gamma$) and the MF set but not the GCF set ($\delta$).

Given a GCF $G$ and a MF $M$, and a scoring function $\sigma$, we want to be able to calculate the $p$-value for the potential link between $G$ and $M$, i.e. the probability that $\sigma(M, G) > s$ under the assumption that $G$ and $M$ are independent. This means that for our purposes, the sizes of $M$ and $G$, $\#M$ and $\#G$, are constants, as well as the total population size, $\#N$.

Assuming that $M$ and $G$ are independent, we have given only $\#M$ and $\#G$, and not $\#(M \cap G)$. Therefore, we have

$$p(\sigma(M,G) = s) = \sum_{o\in\mathbb{N}} p(\#(M\cap G) = o)p(\hat{\sigma}(o) = s) \quad (3)$$

We can therefore calculate the $p$-value as

$$p(\sigma(M,G) > s) = \sum_{s'>s} p(\sigma(M,G) = s') \tag{4}$$

$$= \sum_{s'>s} p(\sigma(M,G) = s' \mid \#M, \#G, \#N) \tag{5}$$

$$= \sum_{s'>s} \sum_{o\in\mathbb{N}} p(\#(M \cap G) = o) p(\hat{\sigma}(o) = s') \tag{6}$$

But since the score is determined completely by $\#M$, $\#G$, $\#(M \cap G)$ and $\#N$, the last term is always either 0 or 1. Furthermore,

$$p(\hat{\sigma}(\#(M \cap G)) = s') = 1 \tag{7}$$

precisely for those values of $\#(M \cap N)$ where

$$\hat{\sigma}(\#(M \cap G)) = s'.$$

Assuming as we are that $M$ and $G$ are independent, the first term, $p(\#(M \cap G) = o)$, can be considered, in terms of a hypergeometric distribution, as the probability of $o$ "successes" in $\#M$ draws from a population of $\#N$, with a total of $\#G$ marked elements, i.e.

$$p(\#(M \cap G) = o) = p(o \mid \#M, \#G, \#N)$$

follows hypergeometric distribution.

The two sums can be taken together as

$$\sum_{o|\hat{\sigma}(o)>s} p(o \mid \#M, \#G, \#N)$$

where $o = \#(M \cap G)$. As we are assuming that $M$ and $G$ are independent, $p(o)$ follows hypergeometric distribution and can be calculated by standard means.

In practice, the calculations only need to be carried out for the values of $o$ which are possible given $\#M$, $\#G$ and $\#N$, i.e. the lower bound for $o$ is $max(0, (\#M + \#G) - \#N)$, and the upper bound is $min(\#M, \#G)$.

#### 2 The NPLinker Framework

NPLinker http://www.github.com/sdrogers/nplinker is intended to address the significant bottleneck that exists in the realization of the potential of genome-led metabolite discovery, namely the slow manual matching of predicted biosynthetic gene clusters (BGCs) with metabolites produced during bacterial culture; linking phenotype to genotype.

The NPLinker tool and its associated web application implement a new data-centric approach to alleviate this linking problem by searching for patterns of strain presence and absence between groups of similar spectra (molecular families; MF) and groups of similar BGCs (gene cluster families; GCF). Searching can be performed based on a number of available analysis methods that can be employed in isolation or combined as required.

NPLinker is implemented as a standard Python package, appropriate for use in scripting or for interactive environments such as a Jupyter notebook.

On top of this package, we have developed a Dockerised NPLinker web application which allows users to run it in a browser and avoid the process of installing and configuring a suitable local Python environment. Interactive HTML widgets (e.g. buttons, tables, sliders) provide a desktop-like interface while a backend Python process responds to the events triggered in the browser.

In order to use the web application, all that is required from the user is a suitable dataset. These may be drawn from the Paired omics Data Platform, from wholly local data, or a combination of the two. In the case of the Paired omics Data Platform the web application is capable of automated downloading and preprocessing, including running BiG-SCAPE if necessary. Output from BiG-SCAPE and other potentially lengthy analysis processes is cached locally to reduce the time required to launch the application after the initial run.

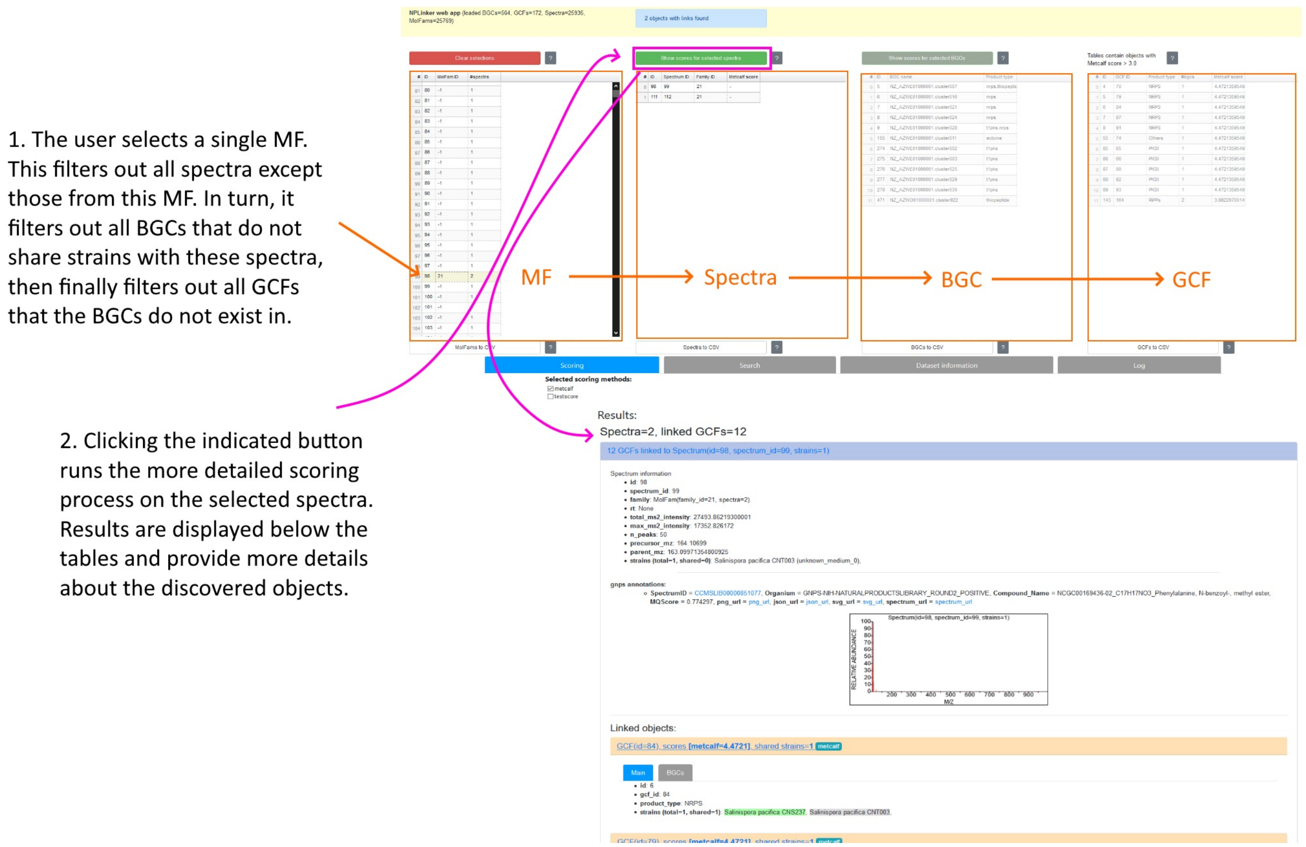


Figure 1: Web application workflow. The user selects an MF from the table on the left, triggering filtering of spectra, BGCs, and GCFs in turn. The button above the spectra table is then clicked to generate results from the enabled scoring methods. The results are then displayed underneath the tables.

The user interface is shown in Figure 1. From left to right the tables contain MFs, spectra, BGCs, and GCFs. The content is determined by an initial run of a standardised correlation scoring on the entire dataset with a user-configurable threshold. This is intended to remove the large number of original objects that are very unlikely to have any significant links. Subsequently when the user makes a selection of one or more objects from any table, all objects that are not linked to the selected object(s) are hidden. For example, selection of a MF will filter out all spectra not contained in that family. This in turn filters out all BGCs that do not share strains with those spectra. Finally, all GCFs that do not contain the BGCs are removed. The filtering operates similarly when other object types are selected as a starting point. This makes it possible for the user to rapidly look up and explore interesting objects in a dataset before running other scoring methods on the filtered results. At any point the user can also click a button to export a CSV file containing the current data displayed in any of the 4 tables for external analysis.

NPLinker contains all functionality described in the paper, as well as an additional scoring system named "Rosetta scoring". This uses a hand-curated translation table between a small subset of GNPS library spectra, and their BGCs as stored in MiBIG. Putative links between a BGC and a spectrum in a data set can be highlighted where the spectrum shows similarity with a GNPS library spectrum in the Rosetta set and the BGC has homology to the corresponding MiBIG entry.

### 3 NPLinker documentation

All Python source code for NPLinker is hosted on GitHub at https://github.com/sdrogers/nplinker.

An associated wiki at https://github.com/sdrogers/nplinker/wiki contains detailed instructions for installing and running the web application described above. The NPLinker web application Docker image is hosted on DockerHub.

API documentation for the NPLinker framework underlying the web application can be viewed at nplinker.readthedocs.io and is automatically updated when the source code changes. The GitHub repository also contains a heavilycommented Jupyter notebook which walks through the process of loading a dataset and using the NPLinker API to explore it and search for links.

### 4 Product types of verified links

| Gross | size (nt) | type | no. genes |
|---|---|---|---|
| BGC0000632 | 16122 | Terpene, Saccharide | 13 |
| BGC0001381 | 210303 | Polyketide | 102 |
| BGC0001842 | 42814 | NRP (other lipopeptide) | 7 |
| BGC0000463 | 43207 | NRP (other lipopeptide) | 4 |
| BGC0001116 | 57999 | NRP, Polyketide | 18 |
| BGC0000399 | 47020 | NRP (cyclic depsipeptide) | 17 |
| BGC0000296 | 66086 | NRP | 28 |
| BGC0001298 | 41001 | Polyketide | 9 |

| Leao | size (nt) | type | no. genes |
|---|---|---|---|
| BGC0001165 | 87427 | NRP, Polyketide (other) | 14 |
| BGC0000962 | 40156 | NRP, Polyketide (other) | 12 |
| BGC0001000 | 41964 | NRP (Other lipopeptide), Polyketide (other) | 8 |
| BGC0001001 | 69900 | NRP, Polyketide (other) | 26 |
| BGC0001560 | 28792 | NRP, Polyketide | 12 |

| Crusemann | size (nt) | type | no. genes |
|---|---|---|---|
| BGC0000827 | 24300 | Alkaloid | 17 |
| BGC0000333 | 47477 | NRP | 23 |
| BGC0000940 | 7328 | Other | 6 |
| BGC0000241 | 62231 | Polyketide (Other), Saccharide (hybrid/tailoring) | 58 |
| BGC0001228 | 37781 | NRP (Cyclic depsipeptide) | 23 |
| BGC0000137 | 91573 | Polyketide | 39 |
| BGC0001830 | 64171 | Polyketide | 23 |

### 5 Raw vs. standardised strain correlation scores

Comparison of raw and standardised strain correlation scores for the Crusemann, Leao and Gross data sets. Black lines represent the scores of validated links.

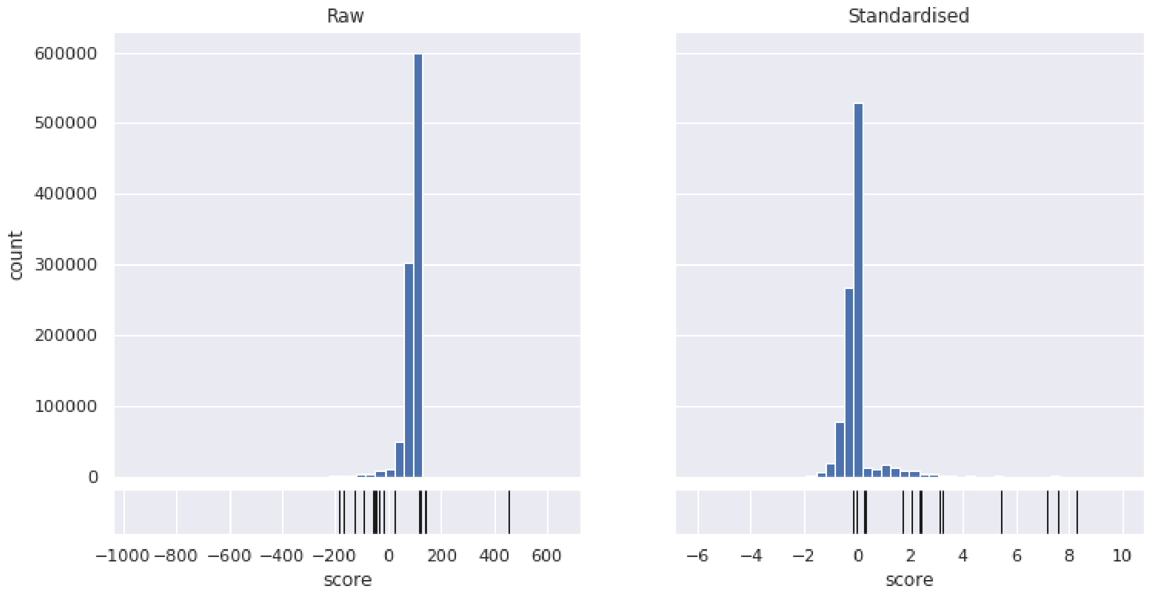

Crüsemann

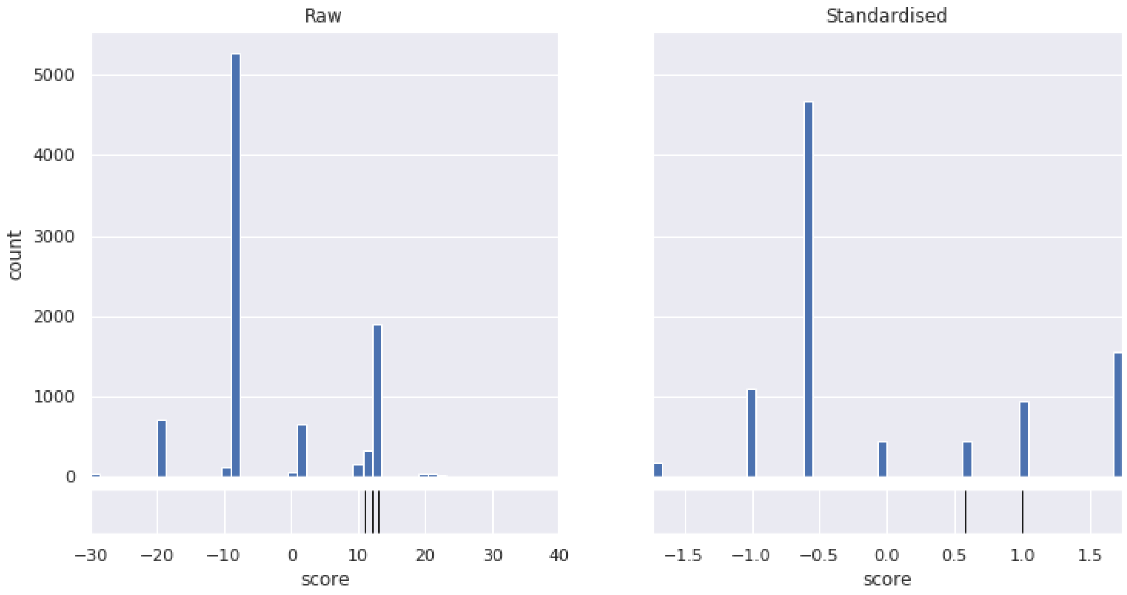


Leao

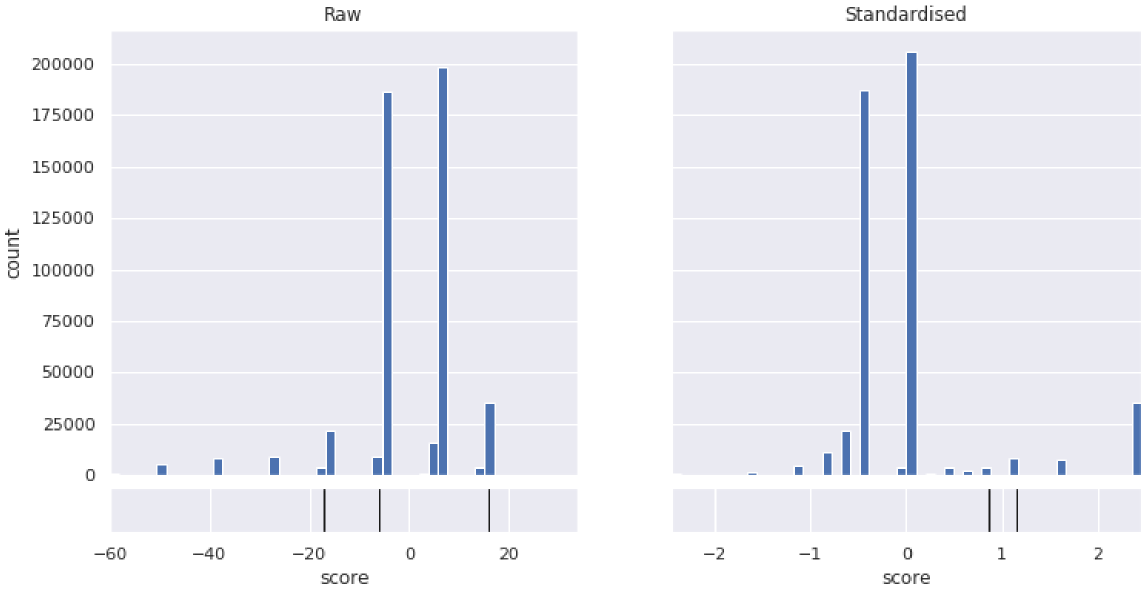


Gross

#### 6 Sizes of data sets

Sizes of the data sets, including how many BGCs could be structurally annotated for IOKR scoring using similarity to MIBiG entries. MIBiG entries are associated with BGCs by running antiSMASH in known cluster blast mode, and any homology detected by antiSMASH is considered valid.

| | Crusemann | Leao | Gross |
|---|---|---|---|
| Total BGCs | 3316 | 147 | 131 |
| BGCs with assigned structure | 2242 | 57 | 83 |
| Total spectra | 6246 | 173 | 9593 |

### 7 IOKR score of validated links

Histograms of IOKR scores for the Crusemann, Leao and Gross data sets. Black lines represent the scores of validated links.

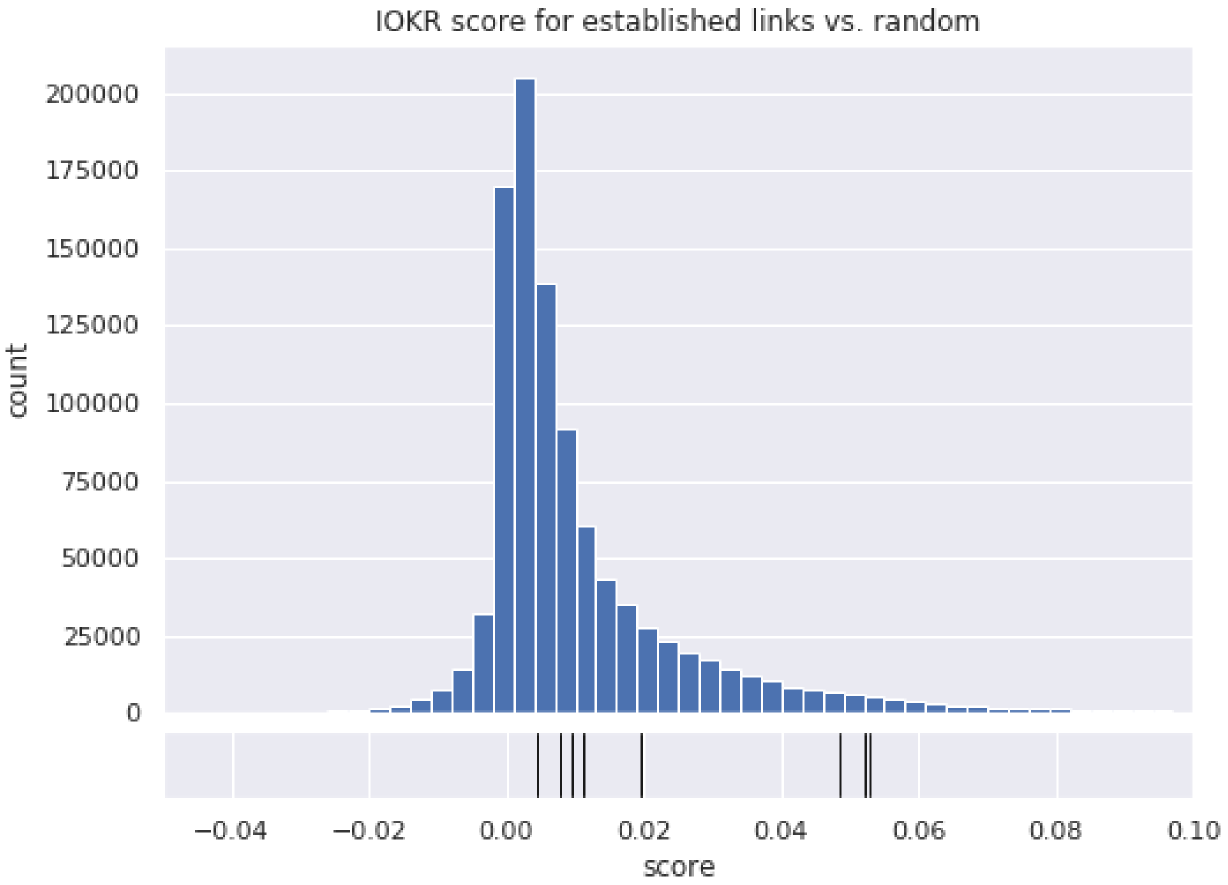


Crusemann

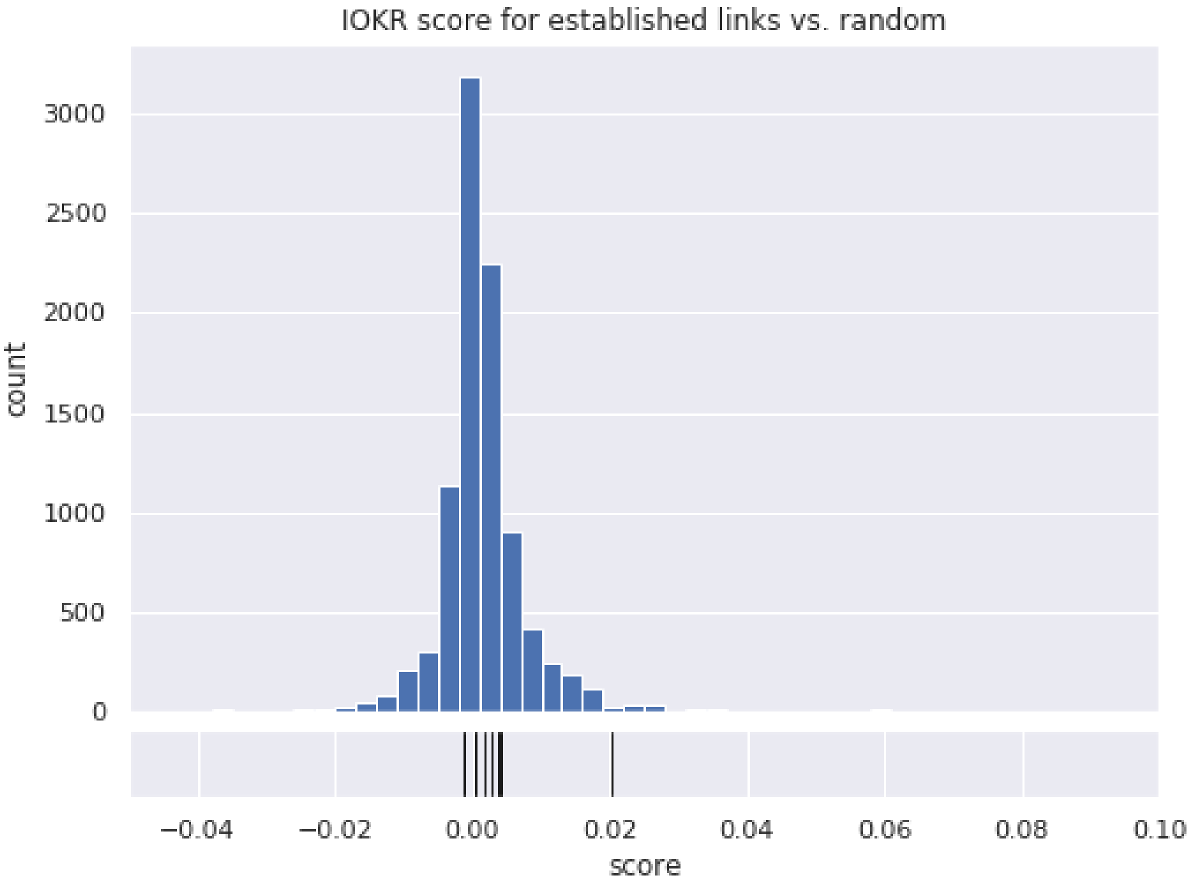
IOKR score for established links vs. random
3000
2500
2000
1500
1000
500
0
count
−0.04
−0.02
0.00
0.02
0.04
0.06
0.08
0.10
score

Leao

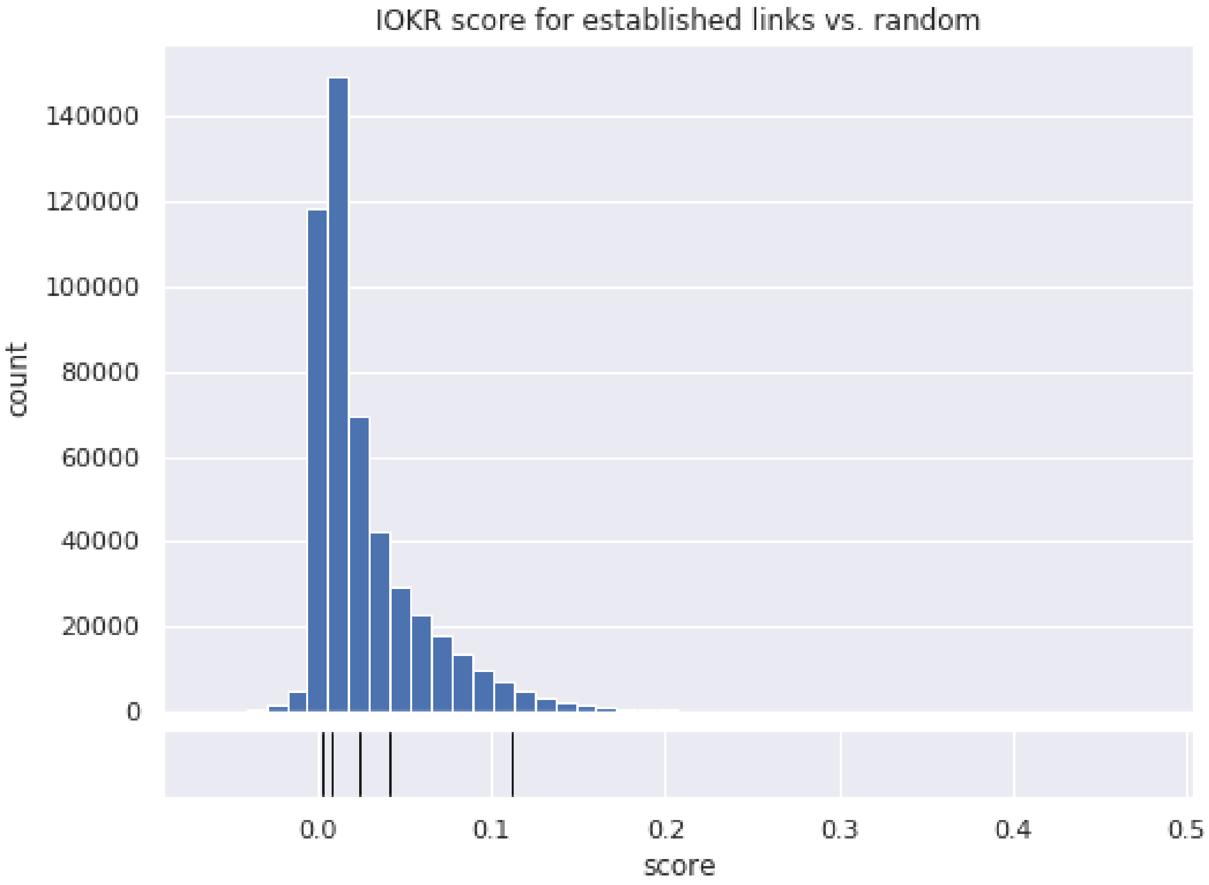


Gross

#### 8 Number of validated links in higher percentiles

Number of links (validated vs. total) for the different scoring functions and data sets. The first two columns describe the unfiltered data, while the others only count links scoring over 95th percentile on either or both scores.

| | all | | > 95% IOKR | | > 95% correlation | | > 95% both | |
|---|---|---|---|---|---|---|---|---|
| | verified | total | verified | total | verified | total | verified | total |
| Gross | 5 | 501886 | 1 | 25095 | 2 | 35333 | 0 | 1537 |
| Leao | 8 | 9342 | 1 | 437 | 6 | 1560 | 1 | 77 |
| Crusemann | 15 | 999362 | 6 | 49970 | 10 | 50224 | 4 | 2517 |
| all | 28 | 1510590 | 8 | 75502 | 18 | 87117 | 5 | 4131 |

| | all | | > 90% IOKR | | > 90% correlation | | > 90% both | |
|---|---|---|---|---|---|---|---|---|
| | verified | total | verified | total | verified | total | verified | total |
| Gross | 5 | 501886 | 1 | 50189 | 4 | 52014 | 1 | 5313 |
| Leao | 8 | 9342 | 1 | 935 | 6 | 1560 | 1 | 147 |
| Crusemann | 15 | 999362 | 6 | 99937 | 13 | 100494 | 5 | 10836 |
| all | 28 | 1510590 | 8 | 151061 | 23 | 154068 | 7 | 16296 |

#### 9 Comparison of score distributions

Comparison of mean scores for all links vs. validated links for the various data sets. The $p$-value is for the null hypothesis that the distributions have identical means.

| Crusemann | Mean score all | Mean score valid | p-value |
|---|---|---|---|
| Raw correlation | 83.5144 | 14.6667 | 0.0001 |
| Standardised correlation | -0.0060 | 3.6717 | 6.8302e-64 |
| IOKR | 0.0105 | 0.0364 | 1.7968e-9 |

| Leao | Mean score all | Mean score valid | p-value |
|---|---|---|---|
| Raw correlation | -1.9843 | 12.625 | 0.0001 |
| Standardised correlation | -0.0218 | 1.4962 | 1.1887e-05 |
| IOKR | 0.0014 | 0.0038 | 0.3922 |

| Gross | Mean score all | Mean score valid | p-value |
|---|---|---|---|
| Raw correlation | -0.7386 | 0.6 | 0.7929 |
| Standardised correlation | 0.0092 | 1.6149 | 6.0056e-06 |
| IOKR | 0.02721 | 0.037020 | 0.5155 |

#### 10 IOKR vs. correlation score

IOKR- ($y$-axis) and strain correlation ($x$-axis) scores for all potential links in the three data sets, with histograms of the scores. Verified links are coloured green on the joint plots. Verified links are concentrated in the upper-right quadrant, i.e. score relatively high on both axes.

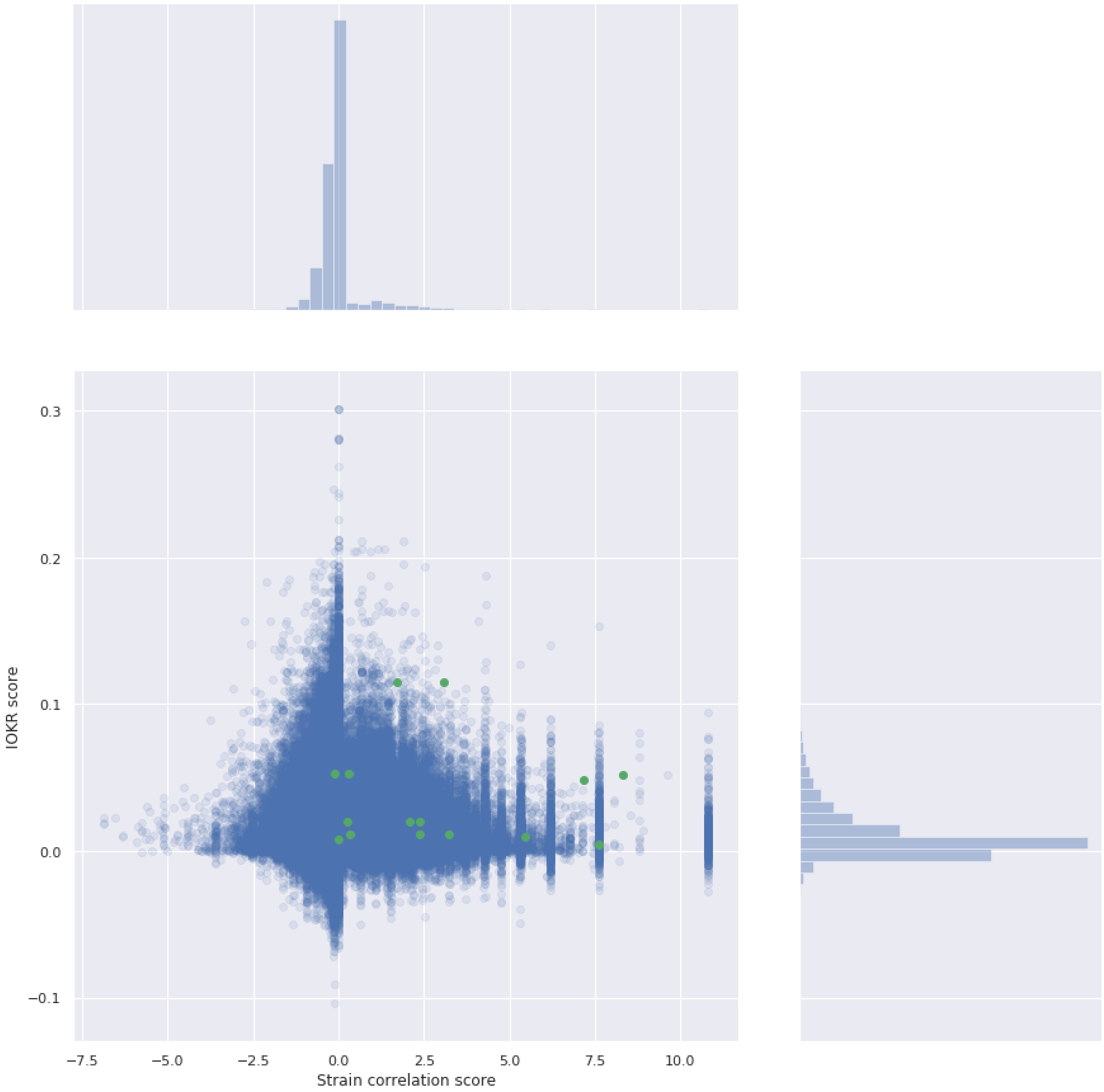

0.3
0.2
0.1
0.0
−0.1
IOKR score
−7.5
−5.0
−2.5
0.0
2.5
5.0
7.5
10.0
Strain correlation score


Crüsemann

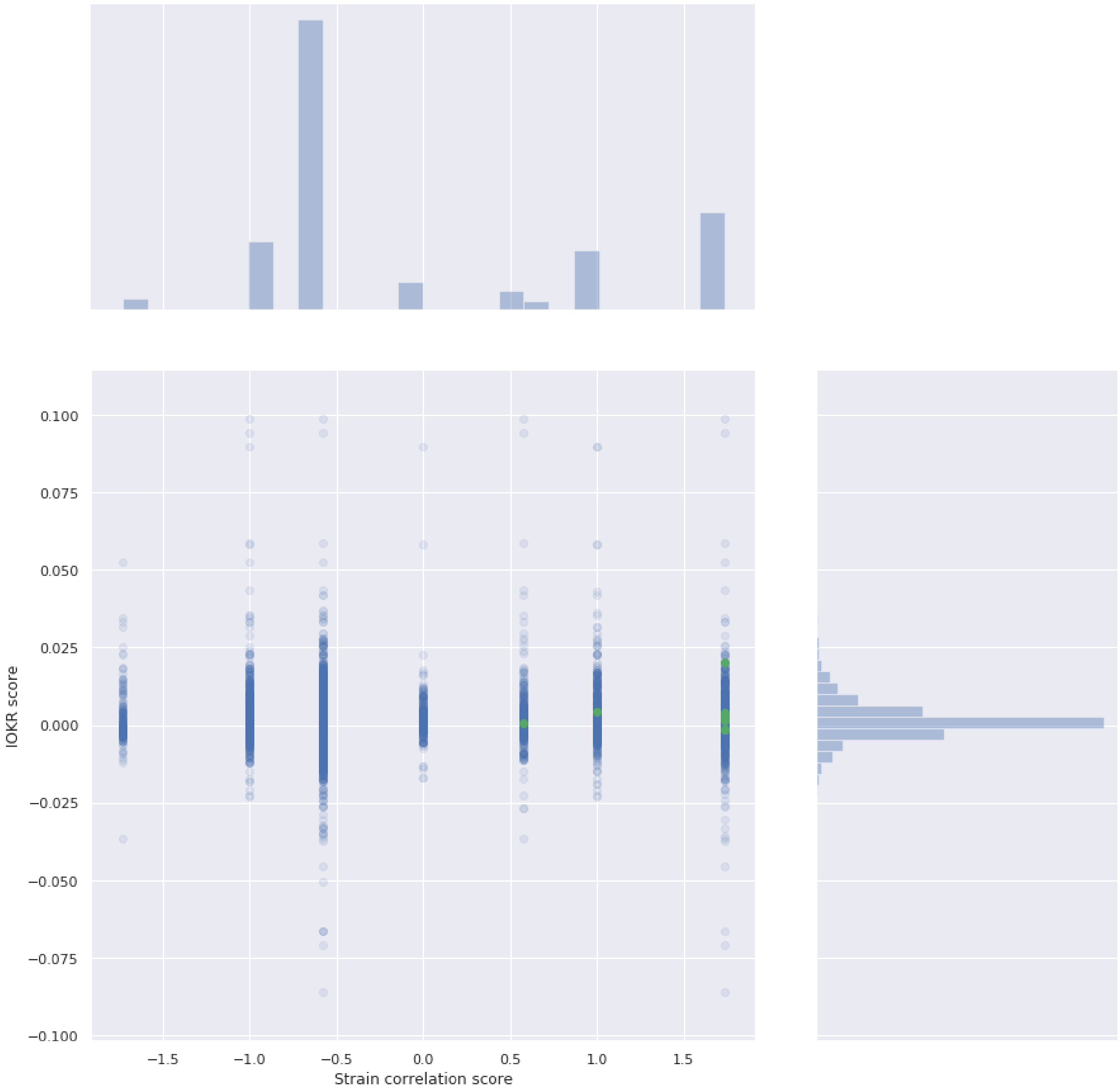
0.100
0.075
0.050
0.025
0.000
−0.025
−0.050
−0.075
−0.100
IOKR score
−1.5
−1.0
−0.5
0.0
0.5
1.0
1.5
Strain correlation score

Leao

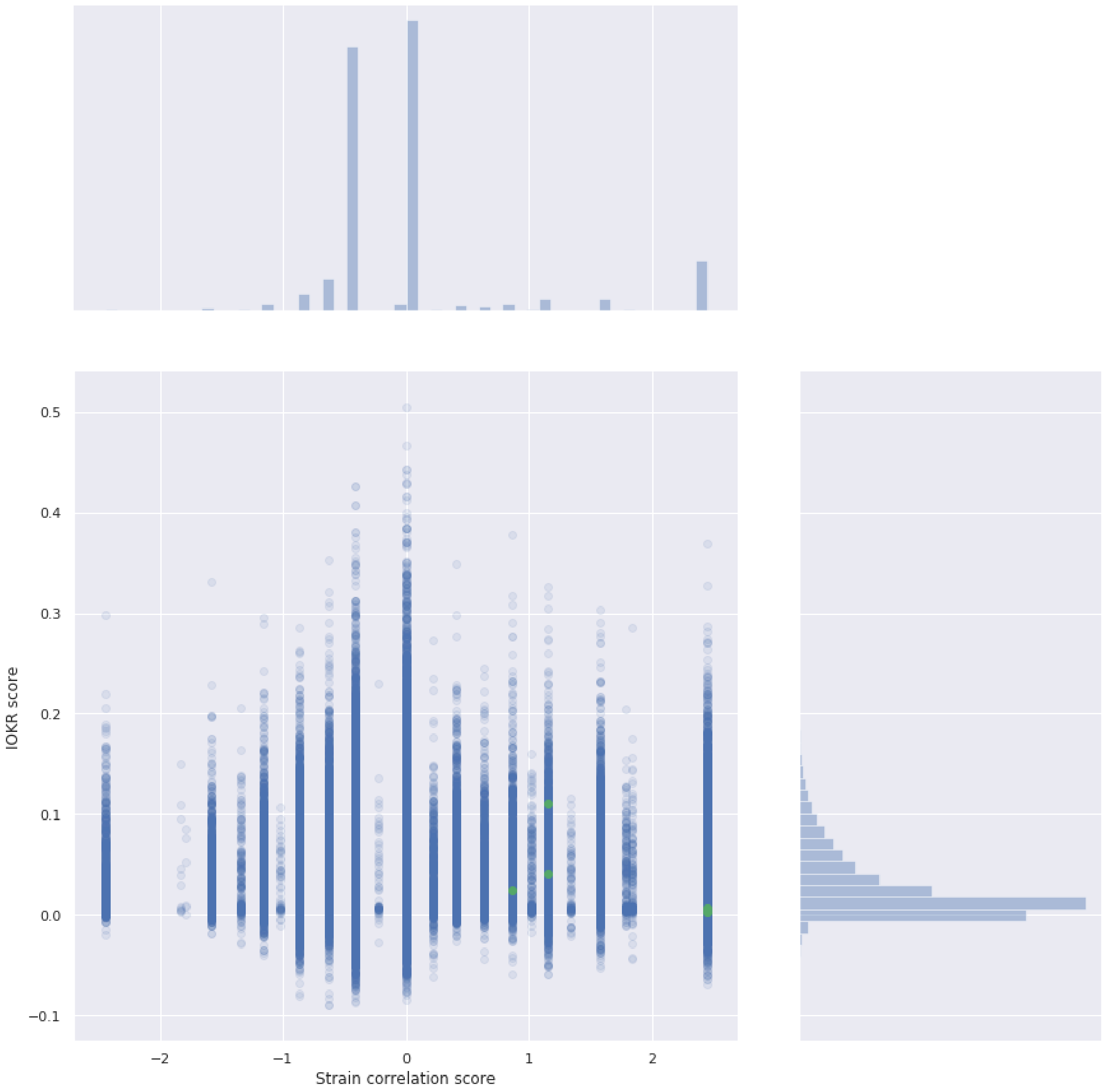


Gross

### 11 Score distributions for a particular BGC

Position of the score for the validated BGC-MF pair (red dot) within the distribution of the scores of the links between that particular BGC and all MFs, for established links (rows). The first three columns show histograms of the raw and standardised versions of the strain correlation score, as well as the IOKR score, for all links including a given BGC, with the score of the correct link indicated. The last column shows the standardised correlation score ($x$-axis) and IOKR score ($y$-axis) for the same links, again with the correct link indicated.

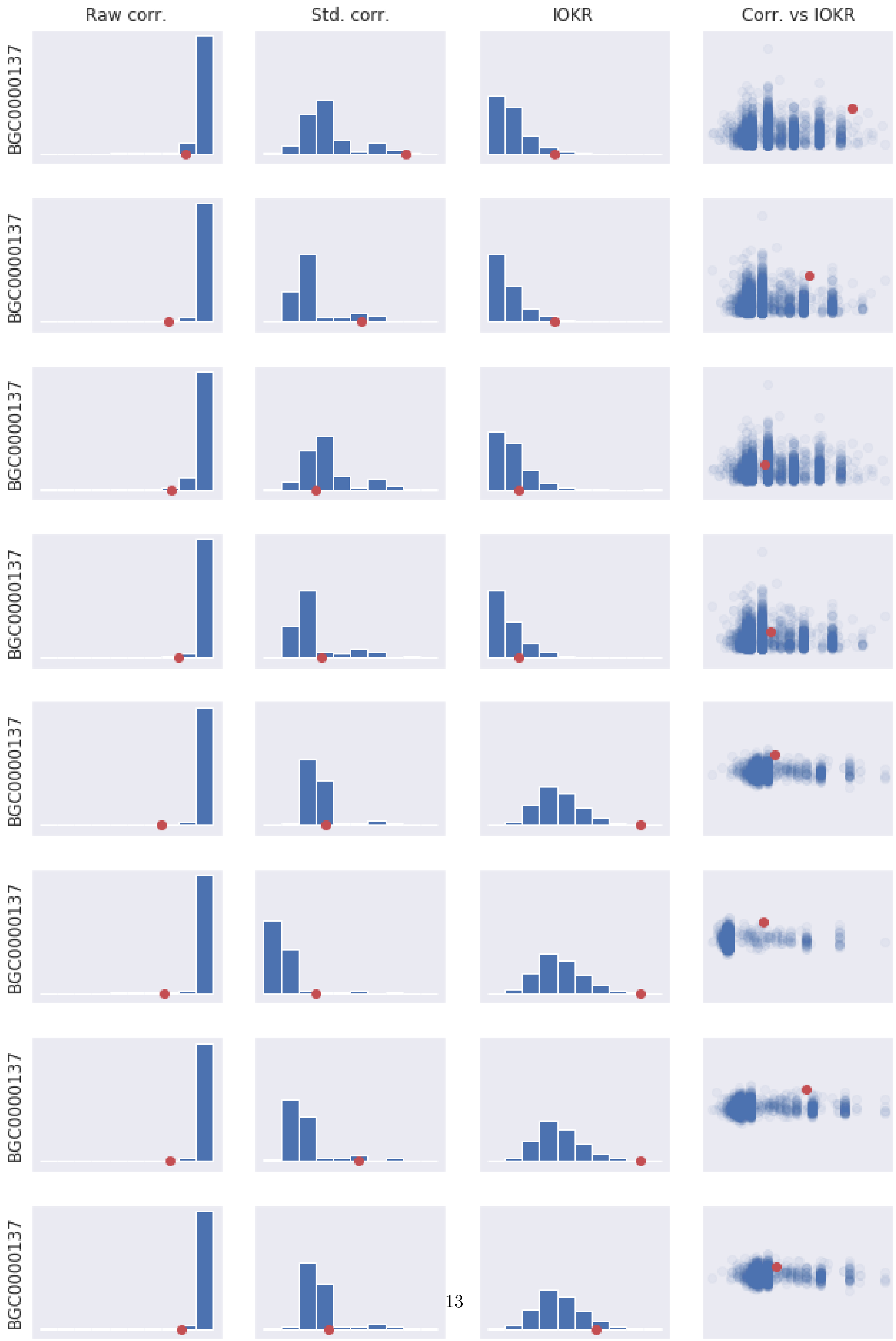
Raw corr.
Std. corr.
IOKR
Corr. vs IOKR
BGC0000137
BGC0000137
BGC0000137
BGC0000137
BGC0000137
BGC0000137
BGC0000137
BGC0000137

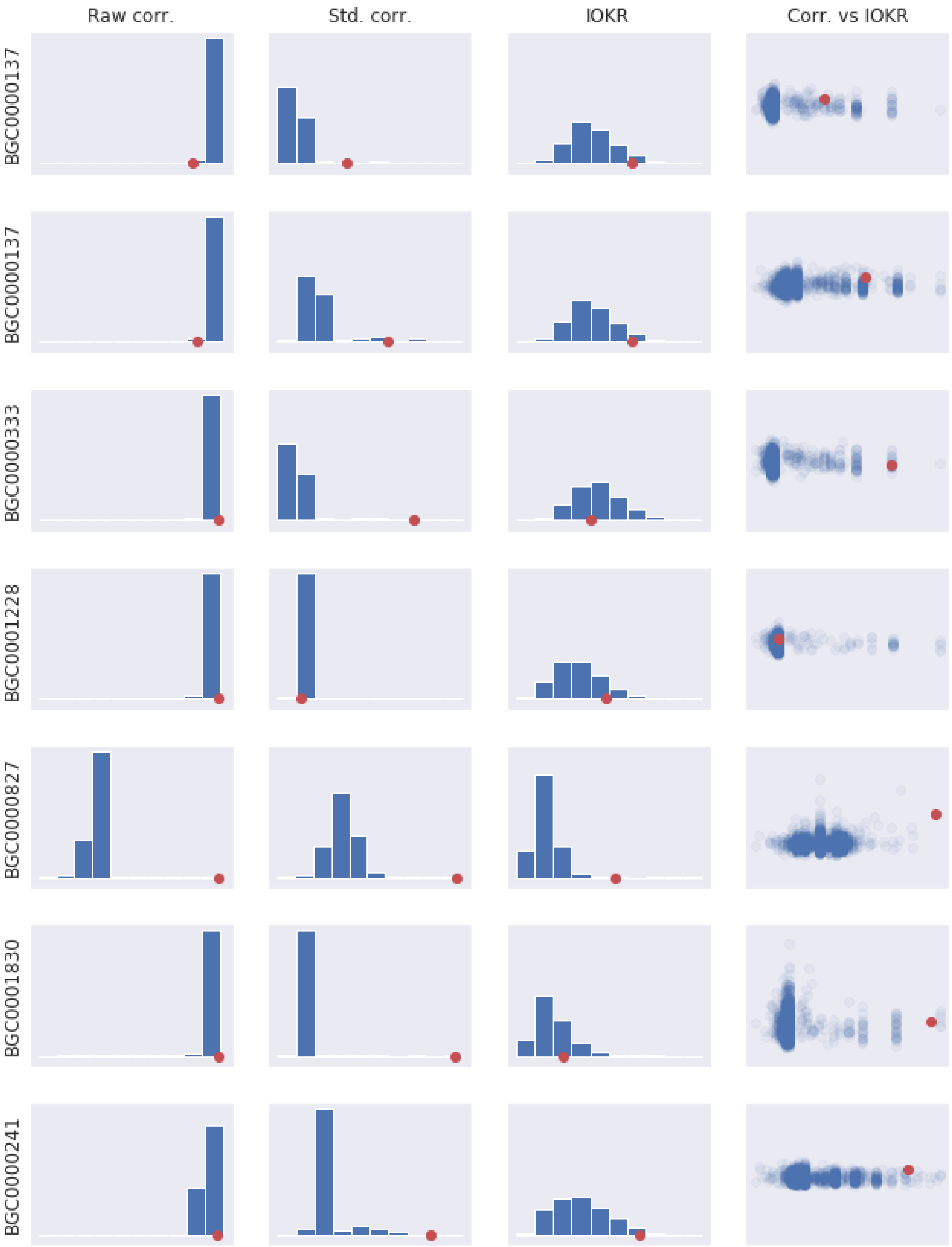
Raw corr.
Std. corr.
IOKR
Corr. vs IOKR
BGC0000137
BGC0000137
BGC0000333
BGC0001228
BGC0000827
BGC0001830
BGC0000241

Crusemann

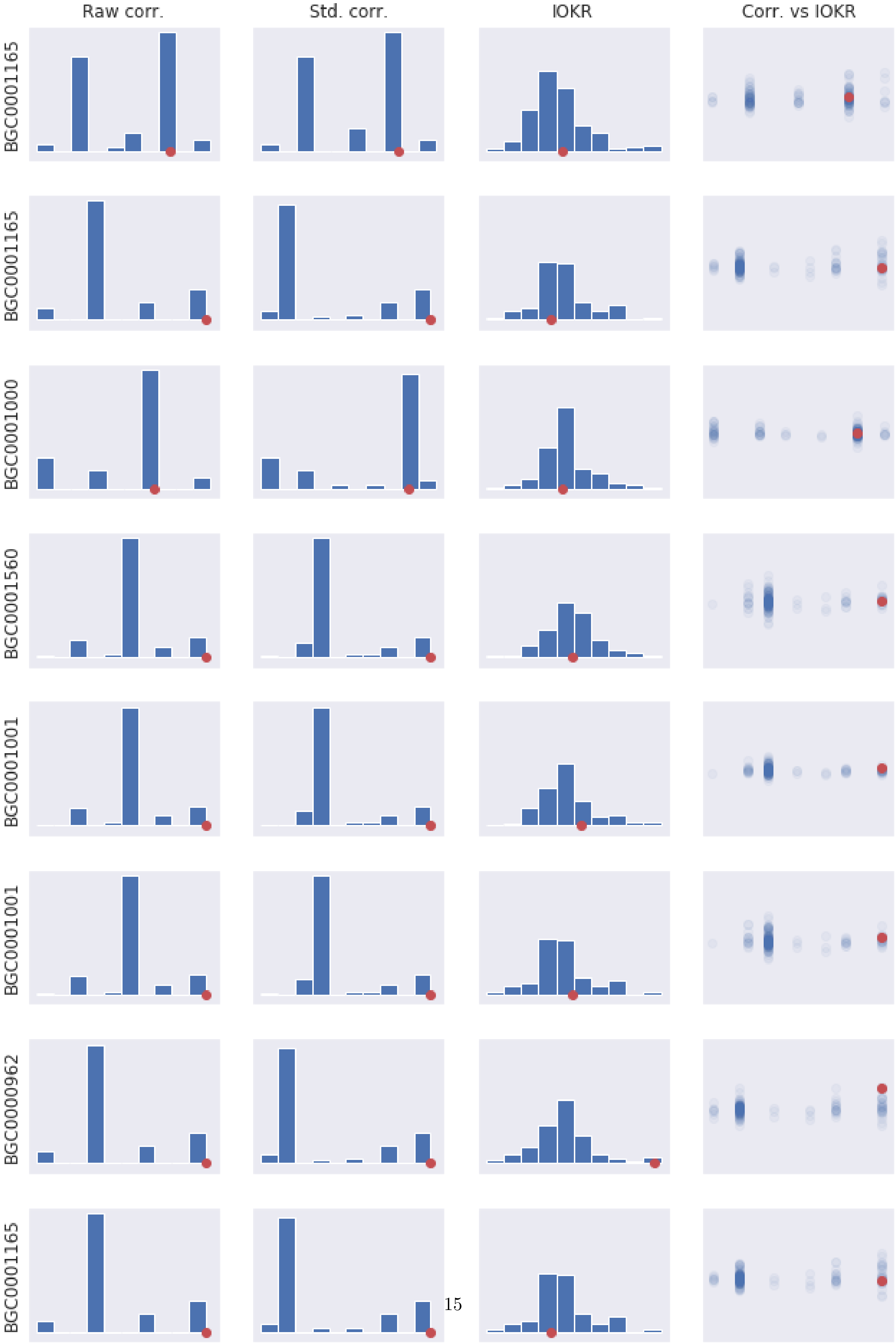

Raw corr.
Std. corr.
IOKR
Corr. vs IOKR
BGC0001165
BGC0001165
BGC0001000
BGC0001560
BGC0001001
BGC0001001
BGC0000962
BGC0001165

Leao

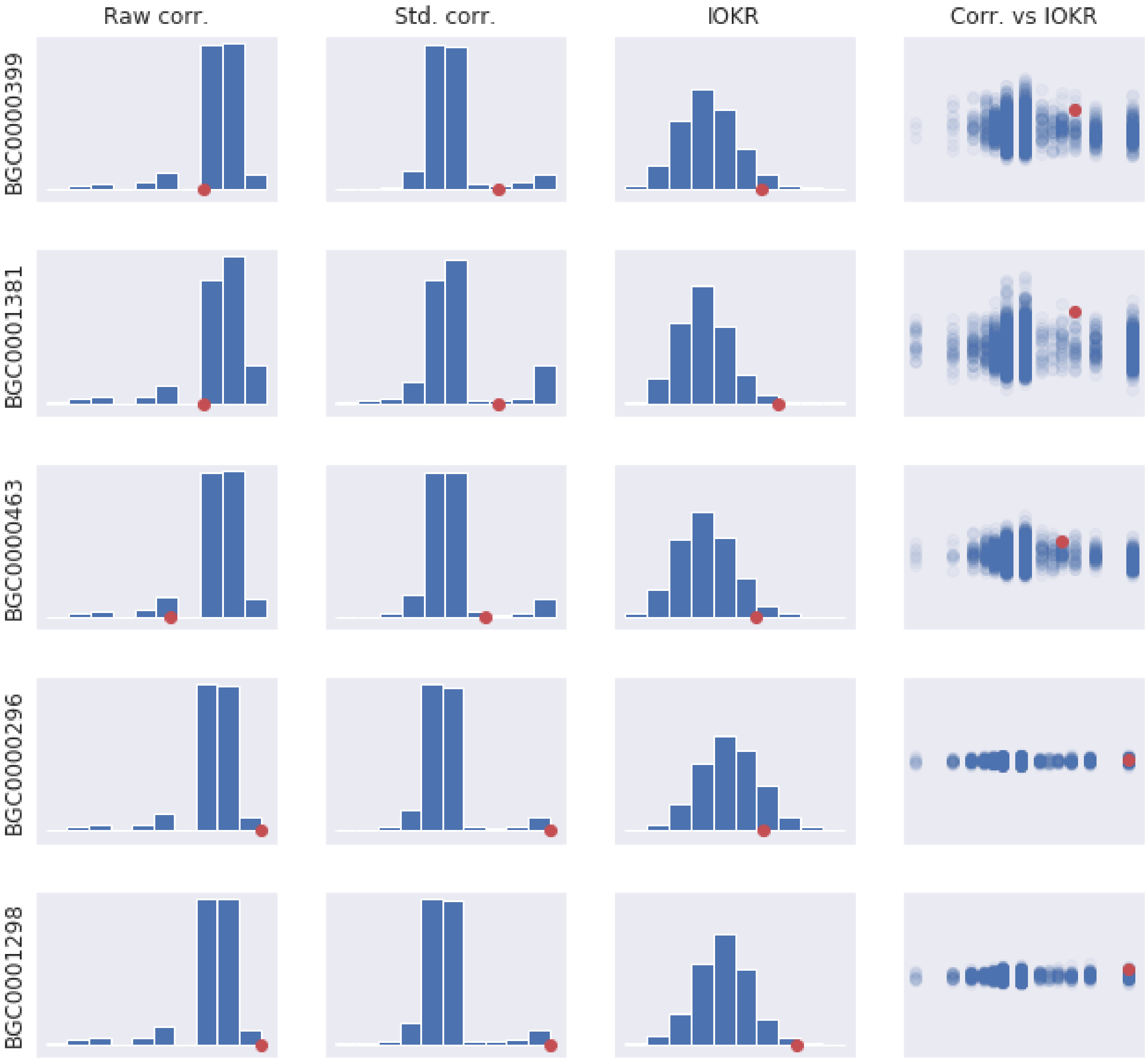


Gross
